# Sex-Dependent Temporal Suppression of Accumbal Dopamine Release During Fentanyl Withdrawal

**DOI:** 10.64898/2026.11.24.768992

**Authors:** Greatness O. Olaitan, Akhabue K. Okojie, Richard Paget-Brown, Parboni Dey, E. Blair Towers, Morgan Gatti, B. Jill Venton, Wendy J. Lynch

## Abstract

**BACKGROUND:** Relapse risk in opioid use disorder persists after acute withdrawal, with women showing greater clinical vulnerability. Reduced accumbal dopamine signaling is implicated in opioid withdrawal, but whether this deficit persists into late fentanyl withdrawal or differs by sex is unknown.

**METHODS:** After 10 days of 24-h/day intermittent-access fentanyl self-administration, female and male Sprague-Dawley rats were tested immediately or during early (days 1– 3) or late (days 13–16) withdrawal, versus saline/drug-naïve controls. We combined in vivo fast-scan cyclic voltammetry (pulse-number, stimulation-frequency, and current-amplitude domains) with Western blotting of five candidate proteins.

**RESULTS:** Control females displayed greater evoked dopamine release than males. Female release stayed elevated immediately after self-administration but declined across withdrawal, abolishing the sex difference; male release was suppressed without further decline after early withdrawal. Versus same-sex controls, late-withdrawal 60-Hz release was 73% lower in females and 46% in males, with pulse and amplitude suppression. Clearance half-time was unchanged; release per pulse decreased. In NAc tissue, females had higher baseline GAPDH, which fentanyl and withdrawal reversed (decreased in females, increased in males). Relative to same-sex saline, β-tubulin increased in females and decreased in males; total DAT tended to decrease; phospho-DARPP-32 and GABA-A α1R were unchanged.

**CONCLUSIONS:** Fentanyl withdrawal abolished the higher baseline accumbal dopamine release of females. Although both sexes reached similar levels in withdrawal, the decline was much larger in females and may contribute to greater craving and withdrawal symptoms. This deficit reflected reduced release per pulse rather than faster clearance and was accompanied by selective, sex-dependent molecular adaptations.

## Introduction

Women with opioid use disorder (OUD) exhibit greater affective and psychiatric burden and are more likely than men with OUD to report using opioids to cope with negative affect (1–5). Females are also more sensitive to drug reinforcement, and women show telescoping, faster progression from first use to treatment (3,6,7). In preclinical fentanyl self-administration models, females show longer-lasting signs of physical dependence, greater vulnerability to relapse, and more rapidly developing withdrawal-related adverse health effects than males (8–10). Despite these pronounced sex differences, their neural basis remains unresolved (1,2). Nucleus accumbens (NAc) core dopamine signaling represents a plausible neural substrate for these effects because it supports reward, motivation, and affective regulation, processes disrupted during opioid withdrawal (11–18). Withdrawal relapse risk is thought to reflect anhedonia linked to reduced dopamine signaling and heightened, glutamate-dependent sensitivity to drug cues (14,18–21). Although opioid withdrawal is generally associated with reduced mesolimbic dopamine activity, evidence specific to fentanyl is limited (18,22–25). In particular, it is unknown whether accumbal dopamine deficits are present immediately after fentanyl self-administration or emerge during withdrawal, whether they persist into late withdrawal, and whether their trajectory differs by sex (18,22,23).

To address these gaps, we used an extended, 24-h/day intermittent-access (IntA) fentanyl self-administration procedure that produces binge-abstinent patterns of intake over the 10-day access period and addiction-relevant withdrawal outcomes following cessation, including physical dependence as defined by withdrawal-induced weight loss, enhanced drug motivation, and incubation of cue-induced drug seeking (1,8,10). Because dopamine-terminal function depends on both release capacity and responsiveness to increasing stimulation demand, we used in vivo fast-scan cyclic voltammetry (FSCV) to assess evoked NAc dopamine release across pulse, frequency, and amplitude stimulation domains. We tested drug-naïve and saline controls, rats immediately after fentanyl self-administration, and rats in early withdrawal (EW; days 1–3) and late withdrawal (LW; days 13–16). We also measured clearance kinetics, release per pulse, and five candidate NAc proteins relevant to dopamine dysfunction, including dopamine transport/signaling (dopamine transporter [DAT], phospho-DARPP-32) (26–29), inhibitory regulation (γ-aminobutyric acid type A receptor α1 subunit [GABA-A α1R]) (30), presynaptic energy metabolism (glyceraldehyde-3-phosphate dehydrogenase [GAPDH]) (31,32), and cytoskeletal/vesicular trafficking (β-tubulin) (33,34). Based on evidence of greater and more persistent withdrawal-related vulnerability in females, we hypothesized that fentanyl withdrawal would produce greater or more persistent accumbal dopamine dysfunction in females than males, resulting in distinct sex-dependent trajectories from the end of self-administration through late withdrawal. We further expected these effects to vary across stimulation domains and to be accompanied by sex- and time-dependent molecular adaptations relevant to NAc dopamine signaling.

## Methods and Materials

### Subjects

All procedures were approved by the University of Virginia IACUC (Protocol #3492) and followed the NIH Guide for the Care and Use of Laboratory Animals. Sexually mature male (N=38) and female (N=32) Sprague-Dawley rats (Charles River, Wilmington, MA), weighing approximately 370 g (male) and 270 g (female) at the start of the study, were individually housed in operant testing chambers (Med Associates, St. Albans, VT, USA) on arrival and remained in these chambers throughout the study.

### Procedure

Surgical, catheter and self-administration procedures were performed as previously described (10). Briefly, to ensure rapid acquisition, rats were pre-trained to lever-press for sucrose pellets (45 mg) in two to three, 24-h/day sessions under a fixed-ratio 1 (FR1) schedule, followed by jugular catheterization and recovery, and fentanyl self-administration training (0.25 µg/kg/infusion; FR1, ≤40 infusions/day). Rats were then given 24-h/day IntA to fentanyl for 10 days, with FR1 access during 5-min trials initiated every 30 min (Figure 1A).

**Figure 1.**
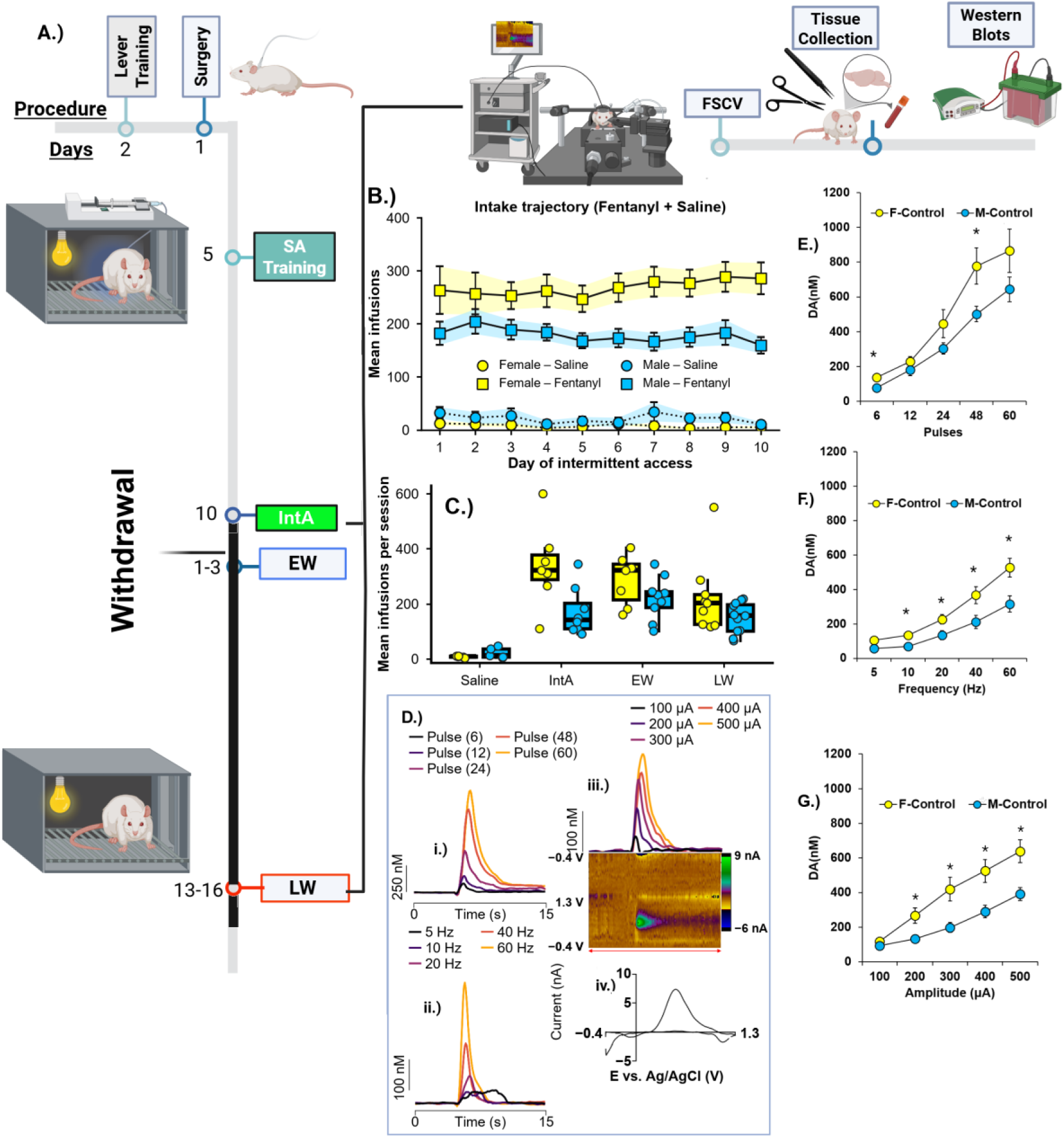
Females take more fentanyl during IntA self-administration than males and exhibit greater baseline evoked dopamine release than males. (A) Experimental timeline. Rats underwent lever-press training, catheter surgery, and fentanyl (0.25 µg/kg/infusion) or saline self-administration training, and once acquired, were given 10 days of 24-h/day access to fentanyl (or saline) under an intermittent-access (IntA) procedure consisting of fixed-ratio 1 access during 5-min trials that initiated every 30 min. Separate cohorts were assessed using FSCV immediately after the final IntA session (IntA), or during early withdrawal (EW; days 1–3), or late withdrawal (LW; days 13–16). Tissue was then collected from the NAc core for Western blotting. Created in BioRender. Lab, L. (2027) https://BioRender.com/g0ifrk9. (B) Mean infusions per session across the 10 IntA days in fentanyl- and saline-self-administering females and males. Fentanyl rats took more infusions than saline rats (Treatment: F(1,57)=40.99, p<0.001). Among fentanyl rats, females self-administered more fentanyl than males (Sex: F(1,49)=11.41, p=0.001) and accumulated more total infusions across the 10 days (2781±274 vs. 1785±145; Welch t(34.0)=3.22, p=0.003); intake did not change significantly across days in either sex (Day: p=0.890; Sex × Day: p=0.120). (C) Mean infusions per session for the rats in each condition (Saline, IntA, EW, LW) by sex; overall mean intake did not differ among the IntA, EW and LW groups within either sex (Holm p≥0.11), although daily intake patterns differed (group × day, p=0.048; Supplementary Table S5). (D) Representative FSCV traces for the pulse (i), frequency (ii) and current (iii) series, with a representative color plot and cyclic voltammogram (iv). (E–G) Evoked dopamine release in Control females and males across the pulse (E), frequency (F) and current (G) series; *p<0.05, females vs. males. Data in B and E–G are mean ± SEM; box plots in C show median and interquartile range with individual animals; female fill = yellow, male fill = sky blue; n per cell in Supplementary Table S2.

Following the 10-day IntA period, rats underwent dopamine assessment and/or tissue collection beginning immediately after the final self-administration session, during EW (days 1–3), or during LW (days 13–16). Assignments were non-systematic and were reviewed to ensure that fentanyl intake and other key behavioral measures were comparable between conditions within sex. Effects were also assessed in additional groups of male (n=5) and female (n=5) saline self-administration control rats, which underwent the same surgical and behavioral procedures but self-administered saline instead of fentanyl, and drug-naïve control rats (n=4 per sex). Because no significant differences were detected between saline and drug-naïve controls within sex across the FSCV stimulation series (Supplementary Figure S1 and Supplementary Table S1), their data were pooled into a single Control group for the FSCV analyses. Initial assignments and final sample sizes by sex, condition, and endpoint are provided in Supplementary Table S2. Final group sizes were as follows: Control n=9 male, n=9 female; IntA n=7 male, n=6 female; EW n=8 male, n=7 female; LW n=7 male, n=8 female.

### Fast-scan cyclic voltammetry

In vivo FSCV was performed under isoflurane anesthesia (2–3%) with body temperature maintained on a heating pad (35). A carbon-fiber microelectrode (7 µm diameter) was lowered into the NAc core (AP +1.2, ML +1.2, DV −6.5 to −7.0 mm) and a bipolar stimulating electrode into the ipsilateral ventral tegmental area (VTA; AP −5.2, ML +1.0, DV −8.5 mm); coordinates follow Paxinos and Watson (36). A triangular waveform (−0.4 to +1.3 V, 400 V/s, 10 Hz) was applied, and each electrode was calibrated post-experimentally in a flow-injection cell against known dopamine standards (0.1-2.0 µM) to convert oxidation current to nM release; dopamine was identified by its background-subtracted voltammogram (37). Phasic and tonic stimulation protocols were adapted from prior in vivo electrochemistry work in the mesolimbic system (37–39). Release was evoked by biphasic VTA stimulation (2 ms/phase) across three stimulation series at one site: pulse number (6-60 pulses, 60 Hz, 300 µA), frequency (5-60 Hz, 24 pulses, 300 µA), and amplitude (100-500 µA, 24 pulses, 60 Hz). Successive stimulations were 5 min apart, and electrode placement in the NAc core was verified histologically. For each series, we computed sub-range averages, area under the curve (AUC; trapezoidal integration), and response slope (linear regression); each was also expressed as a percentage of the sex-matched control mean. Voltammetric traces were extracted and analyzed using the HDCV Voltammetry and Analysis software (40).

Clearance half-time (t1/2), Vmax (Michaelis–Menten fit, Km=0.2 µM), and release per pulse ([DA]p) were derived from the 60-pulse, 60-Hz response (38), as detailed in the Supplementary Methods (per-rat values in Supplementary Table S3).

### Western blot analysis

The five proteins were measured by Western blot in NAc core tissue collected 10 min after voltammetry from a subset of recorded rats (Saline, n = 3–5 per sex; IntA, EW, and LW, n = 3–5 per sex per condition), within a larger companion-study panel. Signal was normalized to total lane protein (Revert 700, LI-COR), expressed as percent of the same-sex saline mean, log10-transformed, and analyzed by two-way ANOVA (Group × Sex) with within-sex Dunnett comparisons against saline (Supplementary Methods).

### Statistical analysis

All statistical analyses were conducted in R (v4.5.2), and FSCV and Western blot data were analyzed blind to group. Daily infusions across the 10-day IntA period were first compared between fentanyl and saline rats using a linear mixed-effects model with fixed effects of Sex, Treatment, Day, and their interactions, and a random intercept for animal. Within fentanyl rats, sex differences and escalation across the 10-day period were evaluated using a Sex × Day linear mixed-effects model, cumulative 10-day intake was compared between sexes using Welch’s t-test, and mean intake across the cohorts subsequently tested at IntA, EW, and LW was compared within each sex using one-way ANOVA.

Evoked dopamine release (AUC, slope, and prespecified low, intermediate, and high stimulation ranges) was analyzed using two-way ANOVA (Condition × Sex across Control, IntA, EW, and LW). Baseline sex differences in Control rats were assessed using two-sample t-tests. To evaluate the magnitude of fentanyl- and withdrawal-induced changes relative to each sex’s own baseline, values were also expressed as a percentage change from the same-sex Control mean and analyzed across IntA, EW, and LW using two-way ANOVA (Condition × Sex), supplemented by one-sample t-tests against 0% change within each sex. Planned pairwise comparisons (Welch’s t-tests) between sexes within each condition and between conditions within each sex, and one-sample t-tests against 0% change, were reported uncorrected unless the Condition × Sex interaction for that endpoint was significant, in which case they were adjusted for family-wise error using the Holm–Bonferroni procedure within each endpoint. Effect sizes are reported as partial eta squared (*η^2^_p_*). Average daily fentanyl intake was tested as a covariate (ANCOVA) and by correlation with dopamine release. Bayes factors for evoked release are given in Supplementary Table S4. Statistical significance was set at *α* = 0.05.

## Results

### Females take more fentanyl during IntA self-administration than males and exhibit greater baseline evoked dopamine release than males

During the 10-day IntA self-administration period, rats self-administering fentanyl exhibited substantially greater intake than saline controls (Treatment: F1,57=40.99, p<0.001; Figure 1B). Among fentanyl rats (n=23 females, 28 males), females self-administered more fentanyl than males (Sex: F1,49=11.41, p=0.001), and intake did not change significantly across days in either sex (Day: F9,434=0.48, p=0.890; Sex × Day: F9,434=1.58, p=0.120). Overall mean intake did not differ among the cohorts later assigned to IntA, EW, and LW within either sex (Holm p≥0.11; Figure 1C), although daily intake patterns differed among these cohorts (Withdrawal group × Day: F18,398=1.64, p=0.048; Supplementary Table S5).

In Control rats, peak dopamine release scaled linearly with pulse number and current amplitude, whereas stimulation frequency elicited steady-state release at low frequencies (5–10 Hz) characteristic of tonic firing and larger, burst-like responses at higher frequencies; background-subtracted cyclic voltammograms displaying characteristic oxidation (+0.6 V) and reduction (−0.2 V) features confirmed the dopamine signal across recordings (Figure 1D.i–iv).

Control females exhibited significantly greater evoked dopamine release than males across much of the stimulation range (Figure 1E–G). Integrated summaries demonstrated significant sex differences across all three stimulation domains: pulse AUC, frequency AUC, and amplitude AUC (Supplementary Table S5). In the pulse domain, the female elevation was significant only at 6 and 48 pulses (p=0.033 and p=0.025; other pulse numbers p≥0.10). In the frequency domain, females displayed higher evoked release across the tonic-to-phasic frequency series, with the clearest divergence emerging at high burst frequencies. In the amplitude domain, sex differences scaled positively with current and were most prominent at intermediate-to-high amplitudes. Withdrawal effects were evaluated on AUC and response slope, followed by prespecified low, intermediate, and high ranges (Figures 2B, 2E–G, 3B, 3F–H, 4B, 4F–J; Supplementary Figure S2 and Supplementary Table S6); the prespecified range and single-level analyses are reported in the Supplementary Results.

**Figure 2.**
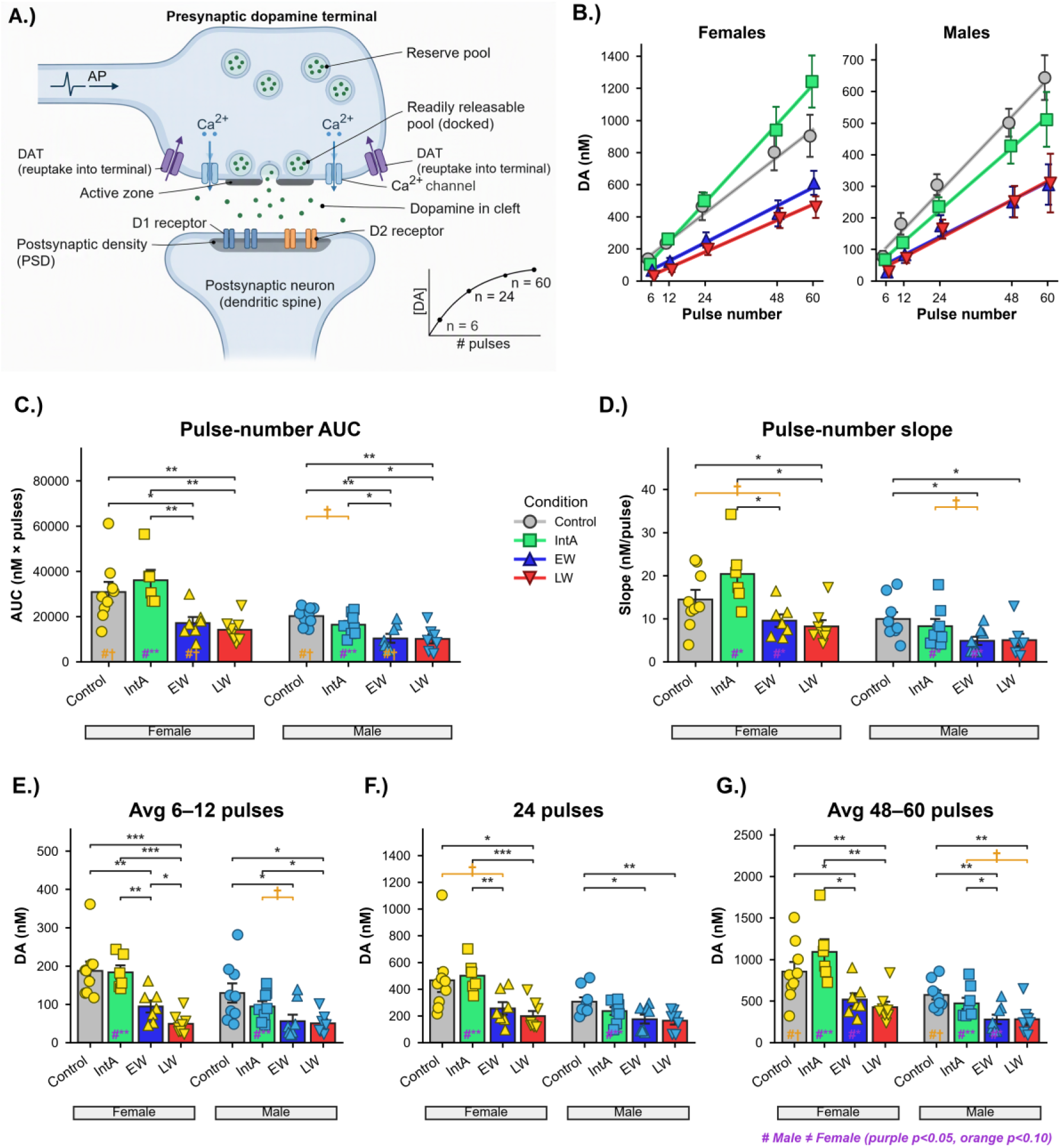
Fentanyl withdrawal produces marked and persistent suppression of pulse-evoked accumbal dopamine release, eliminating the higher female release by late withdrawal. (A) Schematic of a dopamine terminal: action potentials open Ca2+ channels and release dopamine from vesicles of the readily releasable pool docked at the active zone, and DAT takes dopamine back into the presynaptic terminal; released dopamine acts on D1 and D2 receptors at the postsynaptic density (PSD) of a dendritic spine (schematic; D1 and D2 receptors are expressed largely on separate populations of medium spiny neurons). The pulse-number series used 6-60 pulses at 60 Hz and 300 µA. Created in BioRender. Lab, L. (2027) https://BioRender.com/g0ifrk9. (B) Evoked dopamine across the pulse-number series by Sex x Condition (Control, IntA, EW, LW), plotted against pulse number; points are mean +/- SEM and lines are the mean of the per-rat linear fits used for the slope endpoint. (C) Integrated pulse AUC: interaction, F(3,54)=2.75, p=.0518, partial eta-squared=.132; Condition, F(3,54)=13.86, p=8.09×10^-7, partial eta-squared=.435; Sex, F(1,54)=26.09, p=4.36×10^-6, partial eta-squared=.326. Within females, Control versus EW (p=.0220), Control versus LW (p=.0061), IntA versus EW (p=.0069), and IntA versus LW (p=.0033); within males, Control versus EW (p=.0011), Control versus LW (p=.0016), and IntA versus EW (p=.0372). Females exceeded males in IntA (p=.0058); the Control and EW differences were not significant (p=.0507 and p=.0573). (D) Pulse-response slope: interaction, F(3,54)=2.36, p=.082, partial eta-squared=.116; Condition, F(3,54)=8.06, p=1.58×10^-4, partial eta-squared=.309; Sex, F(1,54)=22.22, p=1.75×10^-5, partial eta-squared=.291. Within females, IntA versus EW (p=.0172), Control versus LW (p=.0330), and IntA versus LW (p=.0101) were significant; within males, Control versus EW (p=.0165) and Control versus LW (p=.0375) were significant. Females exceeded males at IntA (p=.0103) and EW (p=.0206). (E–G) Low (6–12 pulses), intermediate (24 pulses), and high (48–60 pulses) pulse-range responses. At 6–12 pulses, interaction, *F*(3,54) = 1.95, *p* = .1328, *η^2^_p_* = .098; Condition, *F*(3,54) = 15.73, *p* = 1.79 × 10^−7^, *η^2^_p_* = .466; Sex, *F*(1,54) = 12.04, *p* = .0010, *η^2^_p_* = .182. At 24 pulses, Condition, *F*(3,54) = 8.78, *p* = 7.73 × 10^−5^, *η^2^_p_* = .328; Sex, *F*(1,54) = 14.50, *p* = .0004, *η^2^_p_* = .212. At 48–60 pulses, interaction, *F*(3,54) = 2.74, *p* = .0523, *η^2^_p_* = .132; Condition, *F*(3,54) = 12.21, *p* = 3.29 × 10^−6^, *η^2^_p_* = .404; Sex, *F*(1,54) = 27.24, *p* = 2.93 × 10^−6^, *η^2^_p_* = .335. Pairwise p values in C and D are uncorrected because the Condition × Sex interaction was not significant for these endpoints (see Methods). Orange † marks p<0.10. In B, y-axis ranges differ between females and males.

### Fentanyl withdrawal persistently suppresses pulse-evoked dopamine release, eliminating the higher female release

Dopamine release was mapped across a 6–60 pulse-number series (60 Hz, 300 µA; Figure 2A). Across the pulse domain, in vivo FSCV revealed marked withdrawal-associated suppression: dopamine release was relatively preserved immediately following the final self-administration session but declined sharply during EW and remained depressed through LW (Figure 2B; Supplementary Table S6).

Integrated pulse AUC (Figure 2C) confirmed this persistent deficit across withdrawal (Condition: *F*_3,54_=13.86, *p*=8.09×10^-7^, *η^2^_p_* =0.435; Sex: *F*_1,54_=26.09, *p*=4.36×10^-6^, *η^2^_p_* =0.326). We also observed a trend for a Sex × Condition interaction (*p* = 0.0518), which appeared to reflect the higher pulse AUC in females under IntA conditions (*p*=0.0058), smaller female elevations under Control conditions and during EW that did not reach significance (*p*=0.0507 and *p*=0.0573, respectively), and convergence of male and female values during LW. Planned comparisons within females demonstrated robust suppression from Control to EW (*p*=0.0220) and LW (*p*=0.0061), as well as marked reductions from IntA (IntA vs. EW: *p*=0.0069; IntA vs. LW: *p*=0.0033). Within males, pulse AUC was also significantly reduced relative to Control at EW (*p*=0.0011) and LW (*p*=0.0016), and dropped between IntA and EW (*p*=0.0372). EW and LW did not differ significantly from each other within either sex, indicating that terminal suppression was established early and persisted without spontaneous recovery through protracted withdrawal.

Withdrawal also blunted overall pulse-to-pulse recruitment efficiency, as measured by response slope (Figure 2D; Condition: *F*_3,54_=8.06, *p*=1.58×10^-4^, *η^2^_p_* =0.309; Sex: *F*_1,54_=22.22, *p*=1.75×10^-5^, *η^2^_p_* =0.291; Condition × Sex: *F*_3,54_=2.36, *p*=0.082, *η^2^_p_* =0.116). Within females, pulse slope was highest during IntA, although this increase relative to Control was not significant (*p*=0.1590), and then dropped significantly from IntA levels during EW (*p*=0.0172; vs. Control: *p*=0.0842) and from both Control and IntA levels during LW (*p*=0.0330 and *p*=0.0101, respectively). In males, pulse slope was highest under Control conditions and was significantly lower during EW (*p*=0.0165) and LW (*p*=0.0375). Between-sex comparisons showed that the female bias observed at baseline was maintained during IntA (*p*=0.0103) and EW (*p*=0.0206), but converged to similarly reduced levels by LW.

Relative to same-sex Controls (Supplementary Figure S3), pulse AUC was reduced by 44.3% at EW and 54.3% at LW in females, and by 49.0% and 50.3% in males; at 6–12 pulses, late-withdrawal release was reduced by 74.0% in females and 61.1% in males.

### The female elevation in burst-frequency dopamine release is lost during withdrawal

Terminals were tested across a 5–60 Hz frequency series (24 pulses, 300 µA; Figure 3A), spanning tonic to phasic firing rates. Withdrawal produced widespread suppression across the frequency series (Figure 3B–C; Supplementary Table S6).

**Figure 3.**
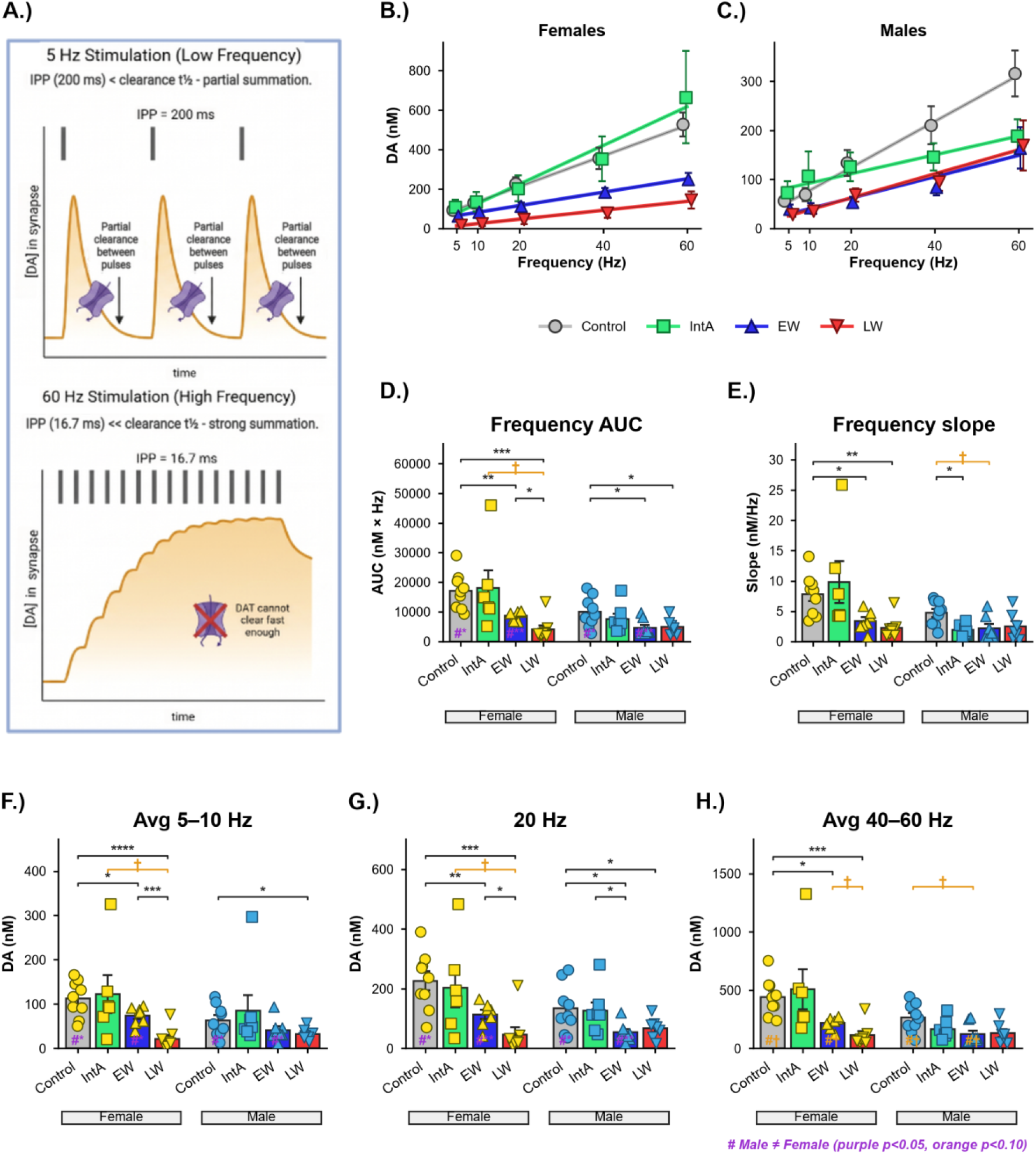
Marked and persistent suppression of burst-frequency-evoked dopamine release across withdrawal, with loss of the female elevation. (A) Frequency-domain schematic (24 pulses, 300 µA): at 5 Hz the 200-ms interpulse interval is shorter than the clearance t½, so release summates partially; at 60 Hz release summates strongly. Created in BioRender. Lab, **L. (2027)** https://BioRender.com/g0ifrk9. (B–C) Evoked dopamine across the frequency series by Sex x Condition, plotted against frequency; points are mean +/- SEM and lines are the mean of the per-rat linear fits used for the slope endpoint. (D) Integrated frequency AUC: interaction, F(3,53)=2.31, p=.0864, partial eta-squared=.116; Condition, F(3,53)=8.73, p=8.42×10^-5, partial eta-squared=.331; Sex, F(1,53)=10.84, p=.0018, partial eta-squared=.170. Within females, Control versus EW (p=.0037), Control versus LW (p=1.37×10^-4), and EW versus LW (p=.0111) were significant; within males, Control versus EW (p=.0150) and Control versus LW (p=.0233) were significant. Females exceeded males in Control (p=.0195) and EW (p=.0035). (E) Frequency-response slope: interaction, F(3,53)=3.83, p=.0147, partial eta-squared=.178; Condition, F(3,53)=5.63, p=.0020, partial eta-squared=.242; Sex, F(1,53)=11.02, p=.0016, partial eta-squared=.172. Within females, Control versus EW (Holm-adjusted p=.0266) and Control versus LW (p=.0060) were significant; within males, Control versus IntA (p=.0201) was significant. Females and males did not differ in any condition (all Holm-adjusted p≥.1477). (F–H) Low (5–10 Hz), intermediate (20 Hz), and high (40–60 Hz) frequency-band responses. At 5–10 Hz, interaction, *F*(3,53) = 1.01, *p* = .3969, *η^2^_p_* = .054; Condition, *F*(3,53) = 6.31, *p* = .0010, *η^2^_p_* = .263; Sex, *F*(1,53) = 4.35, *p* = .0418, *η^2^_p_* = .076. At 20 Hz, interaction, *F*(3,53) = 1.54, *p* = .2141, *η^2^_p_* = .080; Condition, *F*(3,53) = 8.92, *p* = 6.97 × 10^−5^, *η^2^_p_* = .336; Sex, *F*(1,53) = 6.65, *p* = .0127, *η^2^_p_* = .112. At 40–60 Hz, interaction, *F*(3,53) = 3.02, *p* = .0377, *η^2^_p_* = .146; Condition, *F*(3,53) = 7.72, *p* = .0002, *η^2^_p_* = .304; Sex, *F*(1,53) = 11.72, *p* = .0012, *η^2^_p_* = .181. Pairwise p values are uncorrected in D, where the Condition × Sex interaction was not significant, and Holm-adjusted in E, where it was significant. Orange † marks p<0.10. In B and C, y-axis ranges differ between females and males.

Integrated frequency AUC (Figure 3D) did not show a significant Condition × Sex interaction (*F*_3,53_=2.31, *p*=0.0864, *η^2^_p_* =0.116), but showed main effects of Condition (*F*_3,53_=8.73, *p*=8.42×10^-5^, *η^2^_p_* =0.331) and Sex (*F*_1,53_=10.84, *p*=0.0018, *η^2^_p_* =0.170). Between-sex comparisons showed that the female elevation was evident during EW (*p*=0.0035) but not during IntA (*p*=0.1410), and was lost by LW, where female and male values converged. Within females, frequency AUC declined progressively across withdrawal (Control vs. EW: *p*=0.0037; Control vs. LW: *p*=1.37×10^-4^; EW vs. LW: *p*=0.0111). Within males, frequency AUC was also lower during EW and LW than under Control conditions (*p*=0.0150 and *p*=0.0233, respectively).

The overall frequency-response slope (Figure 3E) mirrored this selective loss and showed a significant Condition × Sex interaction (Condition: *F*_3,53_=5.63, *p*=0.0020, *η^2^_p_* =0.242; Sex: *F*_1,53_=11.02, *p*=0.0016, *η^2^_p_* =0.172; Condition × Sex: *F*_3,53_=3.83, *p*=0.0147, *η^2^_p_* =0.178). Within females, frequency slope fell markedly below Control during EW (*p*_Holm_=0.0266) and LW (*p*_Holm_=0.0060), which did not differ from each other. Within males, slope was reduced from Control to IntA (*p*_Holm_=0.0201) and remained numerically below Control during EW and LW, although these differences were not significant (*p*_Holm_=0.0872 and *p*_Holm_=0.2483). Between-sex comparisons showed that the female elevation was no longer significant at IntA (*p*_Holm_=0.2164) and was eliminated during EW and LW.

Consistent with the loss of the female baseline elevation, analysis relative to same-sex Control showed a greater proportional reduction in females by LW, including at 60 Hz (73% reduction in females vs. 46% in males). Males also showed no recovery by LW, with frequency AUC reduced by 54% during EW and 51% during LW.

### Female amplitude-dependent recruitment is relatively preserved before declining by late withdrawal

Dopamine release was measured across a 100–500 µA current series (24 pulses, 60 Hz; Figure 4A). Across the amplitude range, withdrawal produced marked suppression of evoked release (Figure 4B–C; Supplementary Table S6).

**Figure 4.**
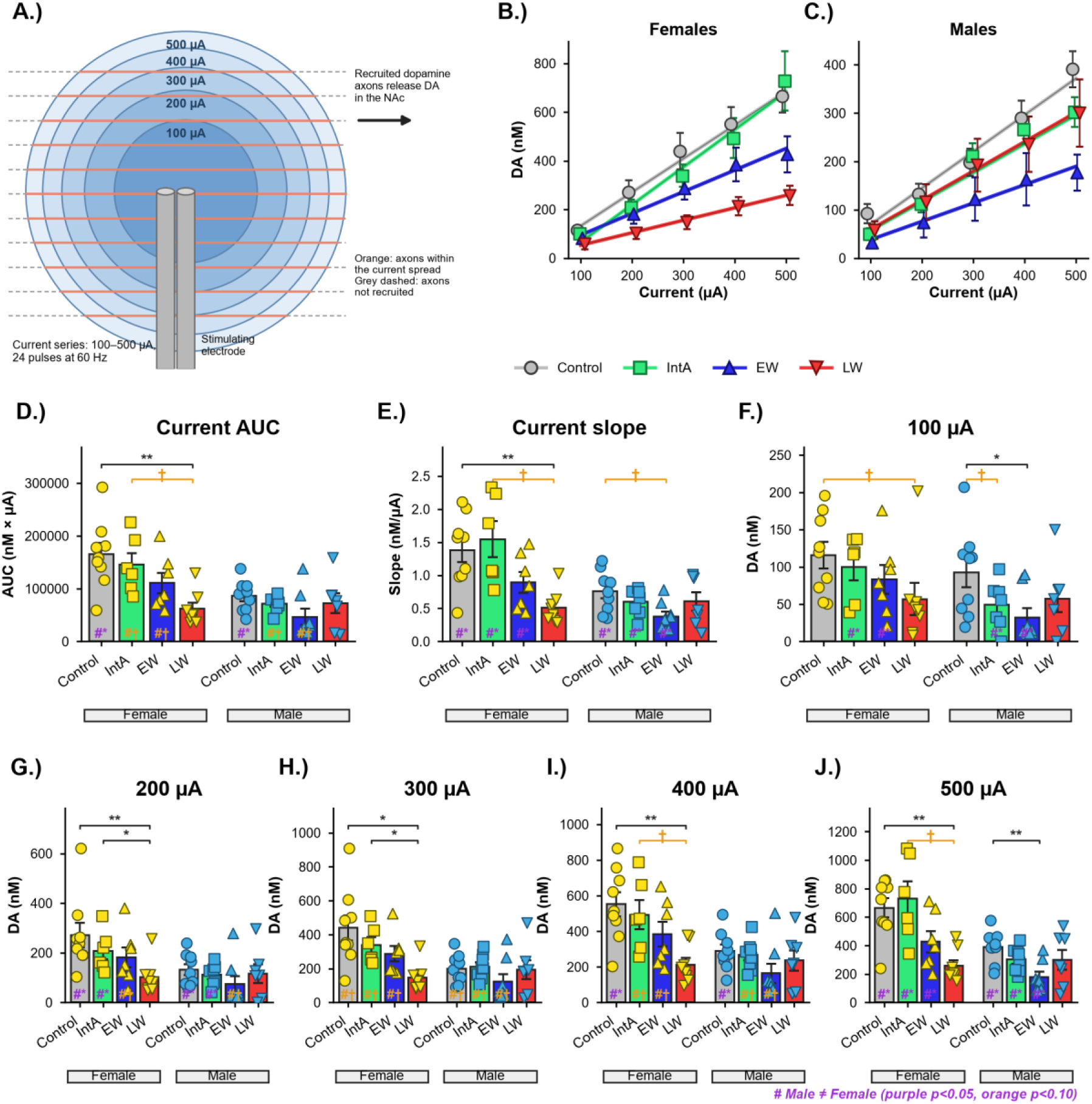
Greater female amplitude-dependent recruitment is lost by late withdrawal. (A) Schematic of current spread at the stimulating electrode: larger currents recruit more dopamine axons (current series, 100-500 µA, 24 pulses at 60 Hz). (B–C) Evoked dopamine across the current series by Sex x Condition, plotted against current; points are mean +/- SEM and lines are the mean of the per-rat linear fits used for the slope endpoint. (D) Integrated amplitude AUC: interaction, F(3,53)=3.33, p=.0262, partial eta-squared=.159; Condition, F(3,53)=6.15, p=.0011, partial eta-squared=.258; Sex, F(1,53)=20.89, p=2.95×10^-5, partial eta-squared=.283. Within females, Control versus LW (Holm-adjusted p=.0064) was significant. Females exceeded males in Control (Holm-adjusted p=.0247). (E) Amplitude-response slope: interaction, F(3,53)=4.36, p=.0081, partial eta-squared=.198; Condition, F(3,53)=7.58, p=2.62×10^-4, partial eta-squared=.300; Sex, F(1,53)=23.52, p=1.12×10^-5, partial eta-squared=.307. Within females, Control versus LW (Holm-adjusted p=.0065) was significant. Females exceeded males in Control (Holm-adjusted p=.0413), IntA (p=.0494), and EW (p=.0494). (F–J) Responses at 100, 200, 300, 400, and 500 µA. At 300 µA, interaction, F(3,54)=3.36, p=.0253, partial eta-squared=.157; Condition, F(3,54)=4.53, p=.0066, partial eta-squared=.201; Sex, F(1,54)=14.57, p=3.50×10^-4, partial eta-squared=.213. Within females, Control versus LW (Holm-adjusted p=.0256) and IntA versus LW (p=.0296) were significant. Females and males did not differ in any condition (all Holm-adjusted p≥.0504). At 500 µA, interaction, F(3,54)=4.95, p=.0041, partial eta-squared=.216; Condition, F(3,54)=9.87, p=2.73×10^-5, partial eta-squared=.354; Sex, F(1,54)=27.71, p=2.50×10^-6, partial eta-squared=.339. Within females, Control versus LW (Holm-adjusted p=.0012) was significant; within males, Control versus EW (p=.0063) was significant. Females exceeded males in Control (Holm-adjusted p=.0156), IntA (p=.0436), and EW (p=.0436). Female amplitude AUC was reduced by 33% in EW and 62.5% in LW relative to the female Control mean. Pairwise p values are Holm-adjusted because the Condition × Sex interaction was significant for each endpoint shown. Bars are mean +/- SEM; points are individual animals; exact n and complete contrasts are in Supplementary Tables S2 and S6. Orange † marks p<0.10. In B and C, y-axis ranges differ between females and males.

Integrated amplitude AUC (Figure 4D) revealed a significant Condition × Sex interaction (*F*_3,53_=3.33, *p*=0.0262, *η^2^_p_* =0.159), alongside main effects of Condition (*F*_3,53_=6.15, *p*=0.0011, *η^2^_p_* =0.258) and Sex (*F*_1,53_=20.89, *p*=2.95×10^-5^, *η^2^_p_* =0.283). The interaction reflected distinct withdrawal trajectories in females and males. Between-sex comparisons revealed that females had significantly higher amplitude AUC than males under Control conditions (*p*_Holm_=0.0247), with smaller differences at IntA and EW that did not reach significance (both *p*_Holm_=0.0564); by LW, this sex difference was abolished as female AUC continued to decline while male AUC no longer differed from Control. Within females, amplitude AUC progressively decreased across withdrawal, falling numerically below Control during EW and declining further by LW, when AUC was significantly lower than Control (*p*_Holm_=0.0064) but not significantly lower than IntA (*p*_Holm_=0.0515) or EW (*p*_Holm_=0.2084). In males, AUC reached its lowest levels during EW, although this reduction relative to Control was not significant (*p*_Holm_=0.2975), and was closer to Control by LW.

Withdrawal also reduced amplitude-response slope (Figure 4E; Condition: *F*_3,53_=7.58, *p*=2.62×10^-4^, *η^2^_p_* =0.300; Sex: *F* =23.52, *p*=1.12×10^-5^, *η^2^_p_* =0.307; Condition × Sex: *F*_3,53_=4.36, *p*=0.0081, *η* ^2^=0.198). Female slopes significantly exceeded male slopes at IntA (*p*_Holm_=0.0494) and EW (*p*_Holm_=0.0494), with sex convergence emerging at LW. Within females, recruitment slope decreased progressively over withdrawal, with a significant decrease from Control during LW (*p*_Holm_=0.0065); decreases from IntA to LW (*p*_Holm_=0.0564), from IntA to EW and from EW to LW were not significant (both *p*_Holm_=0.2371). In males, slope was lowest during EW, but no condition differed significantly from Control (Control vs. EW: *p*_Holm_=0.0640).

Thus, amplitude-dependent recruitment identifies a distinct temporal profile: females retained robust high-current responsiveness immediately after self-administration and through EW, but this capacity collapsed by LW, converging with suppressed male release. Relative to same-sex Control, female amplitude AUC was reduced by 33% during EW and 62.5% during LW, whereas males showed their largest proportional decrease during EW. A similar pattern was observed for amplitude-response slope, with reductions persisting through LW in females but not males. Across all three domains, EW and LW did not differ in males on any of the 18 endpoints (all p≥0.28), whereas female release was lower in LW than EW at low stimulation (5–10 Hz, frequency AUC, 6–12 pulses, 20 Hz; p=0.0006– 0.0365) and was below Control on 17 of 18 endpoints in LW versus 9 of 18 in EW.

### Withdrawal alters dopamine release dynamics and candidate NAc proteins

Clearance half-time of the 60-pulse, 60-Hz response did not differ by Condition, Sex, or interaction (all p≥0.36; Figure 5A–B). Instead, less dopamine was released per pulse during withdrawal ([DA]p: Condition, F3,30=9.11, p=0.0002, ηp2=0.477; Sex, F1,30=4.35, p=0.046, ηp2=0.127; interaction, F3,30=1.84, p=0.161; Figure 5C). In females, [DA]p fell from 40.0 nM at IntA to 15.8 and 8.7 nM during EW and LW (p=0.006 and p=0.005), below Control (p=0.031); in males it was about 60% below Control (p≥0.070). Vmax also differed by Condition (F3,30=6.37, p=0.002) and Sex (F1,30=5.19, p=0.030; Figure 5D): it was highest at IntA and lowest during EW and LW (Control, IntA, EW, LW: females 1.04, 1.88, 0.43, 0.42 µM/s; males 0.48, 0.81, 0.38, 0.31 µM/s), but no contrast with Control was significant (p≥0.075).

**Figure 5.**
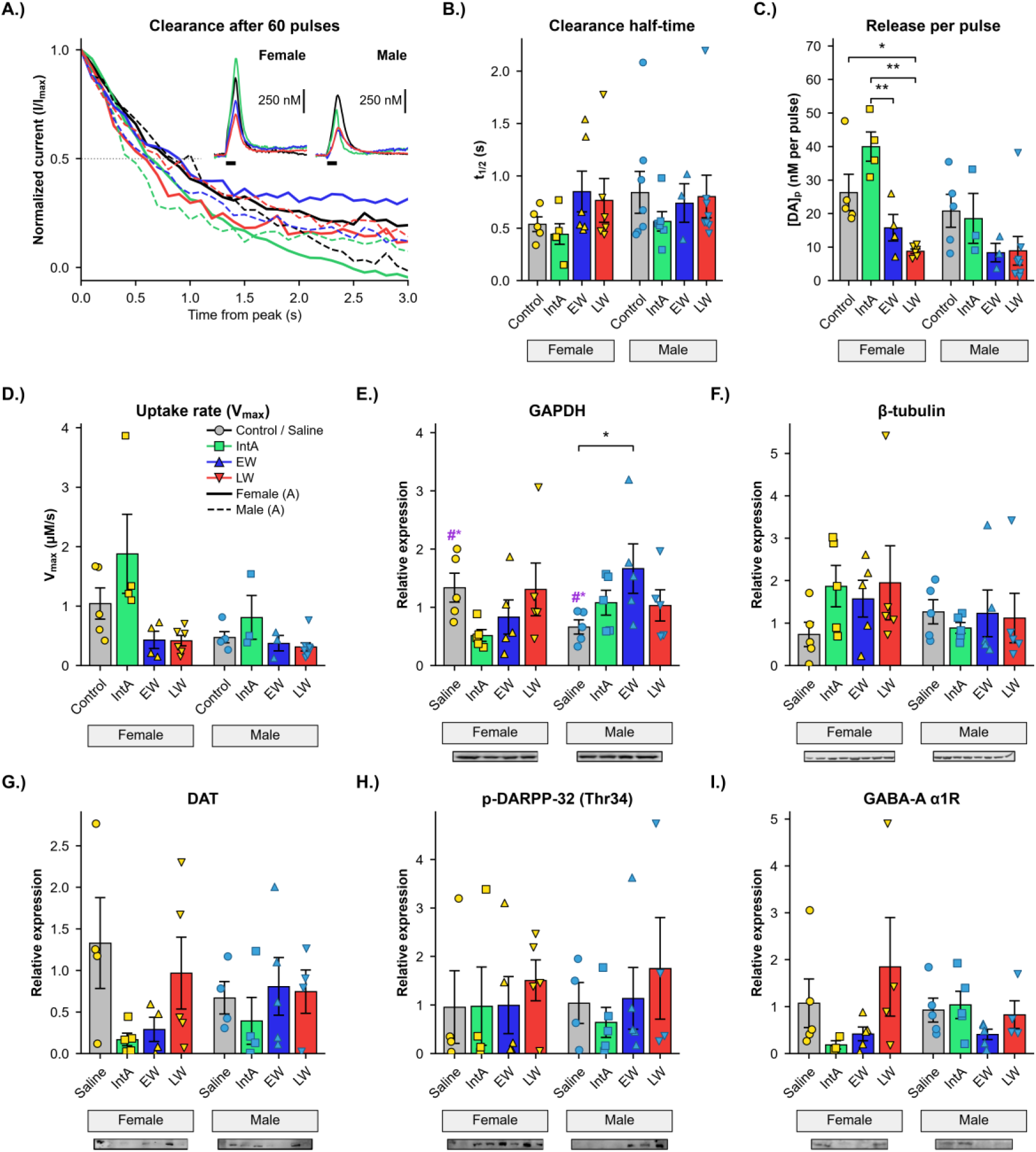
Withdrawal-associated suppression of dopamine release reflects reduced release per pulse rather than faster clearance. (A) Mean peak-normalized clearance after the 60-pulse response in females (solid) and males (dashed); insets show representative dopamine responses to 60 pulses at 60 Hz in females and males from Control, IntA, EW, and LW rats, and the bar under each trace marks the 1-s stimulation. (B) Clearance half-time (t1/2): Condition, F(3,39)=1.10, p=.360, partial eta-squared=.078; Sex, F(1,39)=0.62, p=.437; interaction, F(3,39)=0.39, p=.761. (C) Dopamine released per pulse ([DA]p): Condition, F(3,30)=9.11, p=.0002, partial eta-squared=.477; Sex, F(1,30)=4.35, p=.046, partial eta-squared=.127; interaction, F(3,30)=1.84, p=.161. Within females, IntA versus EW (p=.006), IntA versus LW (p=.005), and Control versus LW (p=.031). (D) Vmax: Condition, F(3,30)=6.37, p=.002, partial eta-squared=.389; Sex, F(1,30)=5.19, p=.030, partial eta-squared=.147; interaction, F(3,30)=1.57, p=.217. (E–I) GAPDH (E), β-tubulin (F), total DAT (G), phospho-DARPP-32 (Thr34) (H), and GABA-A α1R (I). Values are total-protein-normalized signal scaled to the mean of all saline rats (both sexes), so females and males share one scale, with representative blots below each panel; percent of same-sex saline is shown in Supplementary Figure S6. In C, brackets mark planned comparisons between conditions in females (*p<.05, **p<.01). In E, brackets mark Dunnett comparisons against same-sex saline (*p<.05), and # marks the saline female–male difference (two-sample t-test; #* p<.05); female–male differences within fentanyl groups are shown in Supplementary Figure S6. Bar colour and point shape give the group (Control/Saline, grey circle; IntA, green square; EW, blue up-triangle; LW, red down-triangle); yellow points are females and blue points are males. Full membranes are shown in Supplementary Figures S4 and S5. Bars are mean +/- SEM; points are individual animals. Values were log10-transformed for analysis and are plotted untransformed. Group × Sex ANOVA: GAPDH, p = 0.014. DAT: Group trend, p = 0.096. β-tubulin: Sex effect on percent of control, p = 0.020. Phospho-DARPP-32 and GABA-A α1R: all p ≥ 0.07. Within-sex comparisons used Dunnett’s test in two families (vs Saline; EW and LW vs IntA). Between-sex comparisons within each fentanyl group were Holm-adjusted on percent of same-sex saline (Supplementary Figure S6 and Table S9). Baseline sex differences were tested in saline rats with two-sample t-tests.

We next measured five NAc proteins relevant to dopamine dysfunction (Figure 5E–I). In saline rats, GAPDH was approximately two-fold higher in females than males (202%, p=0.034; Figure 5E), but the difference reversed after fentanyl (Group × Sex, F3,32=4.15, p=0.014; Figure 5E). Relative to same-sex saline, GAPDH was 39%, 62%, and 98% in females at IntA, EW, and LW, versus 163%, 252%, and 156% in males (Supplementary Figure S6). The male increase at EW differed significantly from saline (Dunnett p=0.045), and, on percent of same-sex saline, females and males differed at IntA and EW (both Holm p<0.001; Supplementary Figure S6 and Table S9), but not LW. β-tubulin did not differ between sexes at baseline (Figure 5F) but showed divergent relative changes after fentanyl, increasing to 214–266% of same-sex saline in females and decreasing to 70– 97% in males (Sex, F1,32=5.96, p=0.020; Supplementary Figure S6).

DAT did not differ between sexes in saline rats (Figure 5G) but showed a Group trend (F3,27 = 2.34, p = 0.096; Figure 5G), driven by lower levels at IntA in both sexes (females 13%, males 58% of saline) and, in females, at EW (22%), with levels near saline by LW (females 73%, males 111%; Supplementary Figure S6). Phospho-DARPP-32 (Thr34) showed no significant effects (all p ≥ 0.52; Figure 5H). GABA-A α1R showed no significant effects (all p ≥ 0.07; Figure 5I).

## Discussion

Females took more fentanyl than males, and fentanyl withdrawal abolished their higher baseline accumbal dopamine release. Under Control conditions, females showed greater evoked dopamine release than males across pulse, frequency, and amplitude measures, particularly under higher-demand stimulation. This female elevation was partly maintained immediately following fentanyl self-administration but was lost during withdrawal, as dopamine release declined markedly during early withdrawal and declined further by late withdrawal. Males also showed withdrawal-associated suppression, although the timing and magnitude varied across pulse, frequency, and amplitude measures.

This higher release may enhance fentanyl reinforcement and contribute to the greater intake and initial vulnerability of females (3,6,41–43). Intake itself did not predict release, either as a covariate (all p≥0.37) or within any condition (all p≥0.086). Prior work has linked greater female dopamine-release capacity to differences in the readily releasable pool and release probability (44–46). We did not measure estrous-cycle phase or circulating hormones, so the present baseline difference cannot be attributed to a specific hormonal mechanism (45–50). Also, dopamine release was assessed under anesthesia, which preserves the intact mesolimbic pathway but does not establish how the observed deficits affect dopamine signaling during behavior; recordings in awake, behaving animals will therefore be required.

Recognizing this baseline difference is important for interpreting the withdrawal phenotype: females began with greater evoked-release capacity and consequently traversed a larger dynamic range as release declined during withdrawal; because late-withdrawal release was similar in both sexes, greater female withdrawal-related vulnerability may reflect this larger shift rather than the absolute level reached. The suppression was already evident during early withdrawal and remained present during late withdrawal, indicating a persistent rather than transient adaptation (23–25,51). The loss was clearest in the 5–60 Hz frequency domain, which models tonic-to-phasic recruitment relevant to motivational signaling (52–55).

During withdrawal, both sexes were left in a low-dopamine state that persisted in males across pulse and frequency measures and deepened in females. Because accumbal dopamine supports reward and motivation, this deficit may contribute to withdrawal-related anhedonia, a driver of relapse (14,18,23), alongside adaptations in NAc neuron subtypes that drive negative affect in fentanyl abstinence (4). The release deficit did not recover in late withdrawal, when cue-induced seeking incubates in this model (1,10,19), so the dopamine deficit alone is unlikely to explain incubated seeking, which instead reflects glutamatergic plasticity in the NAc (20,21); in the same animals, NAc excitatory-contact markers changed in a sex-dependent manner (56).

Clearance half-time (t1/2) did not shorten, so faster uptake does not explain the deficit; Michaelis–Menten analysis instead attributed the loss of evoked release to less dopamine released per pulse. Surprisingly, the clearest effects were observed for GAPDH and β-tubulin, proteins often used as stable housekeeping controls (57,58), rather than markers more directly related to dopaminergic signaling (DAT (26,27) and pDARPP-32 (28,29)) or its inhibitory regulation (GABA-A α1R (30)). pDARPP-32 and GABA-A α1R showed no overall effects, and although DAT was reduced after IntA, it did not parallel the persistent dopamine-release deficit. GAPDH and β-tubulin instead showed marked, sex-dependent regulation following fentanyl self-administration and withdrawal. GAPDH-driven glycolysis supplies ATP for vesicular glutamate uptake, and GAPDH also acts as an endogenous GABA-A receptor kinase (31,32); its expression decreased in females but increased in males. β-tubulin contributes to microtubule-dependent vesicle trafficking and presynaptic organization (33,34) and showed the opposite pattern, increasing in females and decreasing in males relative to same-sex saline. Because these processes support the energetic and trafficking demands of vesicle filling and release (31–34), their altered regulation could plausibly influence dopamine released per pulse. The opposite responses also raise the possibility that females and males reach a similar functional deficit through different molecular adaptations.

However, we measured total protein abundance rather than activity, post-translational state, localization, or cell-type-specific expression, and their causal relationship to the release deficit remains unknown. Phospho-DARPP-32 was normalized to total lane protein rather than total DARPP-32, in tissue collected after voltammetry. Together, the smaller release per pulse points to changes in vesicular dopamine content, release probability, terminal energetics or trafficking, or presynaptic inhibition; increased dynorphin/κ-opioid receptor signaling and D2-autoreceptor inhibition both reduce release at terminals and are worth testing (59,60).

In conclusion, females had a much larger decrease in dopamine from intermittent access to fentanyl through early and late withdrawal and their dopamine levels were similar to males in late withdrawal. Thus, while both sexes were in a low-dopamine state when relapse risk is high, females experienced a larger drop, which may explain why they show signs of higher craving. Reduced release per pulse, together with sex-dependent changes in GAPDH and β-tubulin, points to presynaptic release mechanisms as targets for restoring dopamine function.

## Supporting information

Main Manuscript

## Data and Code Availability

All raw and processed data supporting the findings of this study — raw FSCV recordings, per-animal Western blot quantifications and paired Revert 700 total-protein images, per-metric summary statistics, and source data underlying every figure — together with the complete R analysis and figure-assembly code (analysis pipeline, statistical models, and Bayesian priors) will be deposited on Zenodo with a citable DOI issued at the time of acceptance and mirrored on GitHub.

## Author Contributions

G.O.O., W.J.L., and B.J.V. conceived and designed the study. G.O.O., A.K.O., R.P.-B., P.D., and M.G. performed the experiments and acquired the data, and E.B.T. recorded behavioral data. G.O.O. and W.J.L. analyzed the data and developed the analysis pipeline. G.O.O., W.J.L., and B.J.V. interpreted the results. G.O.O. drafted the manuscript, and all authors critically revised it and approved the final version.

## Acknowledgments

The authors thank the members of the Venton and Lynch laboratories for technical assistance and discussion. This work was supported by the National Institutes of Health (R01DA052893); the funder had no role in study design, data collection and analysis, interpretation, or the decision to submit. Other measures from the same animals are reported in a companion manuscript, and no dataset appears in both.

## References

1. Towers EB, Setaro B, Lynch WJ (2022): Sex- and Dose-Dependent Differences in the Development of an Addiction-Like Phenotype Following Extended-Access Fentanyl Self-Administration. Front Pharmacol 13: 841873.

2. Davis JP, Eddie D, Prindle J, Dworkin ER, Christie NC, Saba S, et al. (2021): Sex differences in factors predicting post-treatment opioid use. Addiction 116: 2116–2126.

3. Lynch WJ (2006): Sex differences in vulnerability to drug self-administration. Exp Clin Psychopharmacol 14: 34–41.

4. Fox ME, Wulff AB, Franco D, Choi EY, Calarco CA, Engeln M, et al. (2023): Adaptations in Nucleus Accumbens Neuron Subtypes Mediate Negative Affective Behaviors in Fentanyl Abstinence. Biol Psychiatry 93: 489–501.

5. Monroe SC, Radke AK (2023): Opioid withdrawal: role in addiction and neural mechanisms. Psychopharmacology (Berl) 240: 1417–1433.

6. Becker JB, Hu M (2008): Sex differences in drug abuse. Front Neuroendocrinol 29: 36–47.

7. Hernandez-Avila CA, Rounsaville BJ, Kranzler HR (2004): Opioid-, cannabis- and alcohol-dependent women show more rapid progression to substance abuse treatment. Drug Alcohol Depend 74: 265–272.

8. Towers EB, Hsu KA, Qillawala EI, Fraser SD, Lynch WJ (2024): Sex Differences in the Development of an Opioid Addiction–Like Phenotype: A Focus on the Telescoping Effect. Biol Psychiatry Glob Open Sci 4: 100373.

9. Towers EB, Williams IL, Qillawala EI, Rissman EF, Lynch WJ (2023): Sex/Gender Differences in the Time-Course for the Development of Substance Use Disorder: A Focus on the Telescoping Effect. Pharmacol Rev 75: 217–249.

10. Bakhti-Suroosh A, Towers EB, Lynch WJ (2021): A buprenorphine-validated rat model of opioid use disorder optimized to study sex differences in vulnerability to relapse. Psychopharmacology (Berl) 238: 1029–1046.

11. Wilson N, Kariisa M, Seth P, Smith H, Davis NL (2020): Drug and Opioid-Involved Overdose Deaths — United States, 2017–2018. MMWR Morb Mortal Wkly Rep 69: 290– 297.

12. Volkow ND (2021): The epidemic of fentanyl misuse and overdoses: challenges and strategies. World Psychiatry 20: 195–196.

13. Comer SD, Cahill CM (2019): Fentanyl: Receptor pharmacology, abuse potential, and implications for treatment. Neurosci Biobehav Rev 106: 49–57.

14. Koob GF, Le Moal M (2001): Drug Addiction, Dysregulation of Reward, and Allostasis. Neuropsychopharmacology 24: 97–129.

15. Volkow ND, Koob GF, McLellan AT (2016): Neurobiologic Advances from the Brain Disease Model of Addiction. N Engl J Med 374: 363–371.

16. Nestler EJ (2005): Is there a common molecular pathway for addiction? Nat Neurosci 8: 1445–1449.

17. Wise RA (2004): Dopamine, learning and motivation. Nat Rev Neurosci 5: 483–494.

18. Diana M (2011): The Dopamine Hypothesis of Drug Addiction and Its Potential Therapeutic Value. Front Psychiatry 2: 64.

19. Grimm JW, Hope BT, Wise RA, Shaham Y (2001): Incubation of cocaine craving after withdrawal. Nature 412: 141–142.

20. Shen H, Moussawi K, Zhou W, Toda S, Kalivas PW (2011): Heroin relapse requires long-term potentiation-like plasticity mediated by NMDA2b-containing receptors. Proc Natl Acad Sci U S A 108: 19407–19412.

21. Doyle SE, Ramôa C, Garber G, Newman J, Toor Z, Lynch WJ (2014): A shift in the role of glutamatergic signaling in the nucleus accumbens core with the development of an addicted phenotype. Biol Psychiatry 76: 810–815.

22. Radke AK, Gewirtz JC (2012): Increased Dopamine Receptor Activity in the Nucleus Accumbens Shell Ameliorates Anxiety during Drug Withdrawal. Neuropsychopharmacology 37: 2405–2415.

23. Diana M, Pistis M, Muntoni A, Gessa G (1995): Profound decrease of mesolimbic dopaminergic neuronal activity in morphine withdrawn rats. J Pharmacol Exp Ther 272: 781–785.

24. Pothos E, Rada P, Mark GP, Hoebel BG (1991): Dopamine microdialysis in the nucleus accumbens during acute and chronic morphine, naloxone-precipitated withdrawal and clonidine treatment. Brain Res 566: 348–350.

25. Acquas E, Di Chiara G (1992): Depression of Mesolimbic Dopamine Transmission and Sensitization to Morphine During Opiate Abstinence. J Neurochem 58: 1620–1625.

26. Vaughan RA, Foster JD (2013): Mechanisms of dopamine transporter regulation in normal and disease states. Trends Pharmacol Sci 34: 489–496.

27. Bu M, Farrer MJ, Khoshbouei H (2021): Dynamic control of the dopamine transporter in neurotransmission and homeostasis. NPJ Parkinsons Dis 7: 22.

28. Yger M, Girault JA (2011): DARPP-32, jack of all trades… master of which? Front Behav Neurosci 5: 56.

29. Nishi A, Kuroiwa M, Shuto T (2011): Mechanisms for the Modulation of Dopamine D1 Receptor Signaling in Striatal Neurons. Front Neuroanat 5: 43.

30. Xi ZX, Stein EA (1998): Nucleus accumbens dopamine release modulation by mesolimbic GABAA receptors—an in vivo electrochemical study. Brain Res 798: 156–165.

31. Laschet JJ, Minier F, Kurcewicz I, Bureau MH, Trottier S, Jeanneteau F, et al. (2004): Glyceraldehyde-3-phosphate dehydrogenase is a GABAA receptor kinase linking glycolysis to neuronal inhibition. J Neurosci 24: 7614–7622.

32. Ikemoto A, Bole DG, Ueda T (2003): Glycolysis and Glutamate Accumulation into Synaptic Vesicles. J Biol Chem 278: 5929–5940.

33. Waites C, Qu X, Bartolini F (2021): The synaptic life of microtubules. Curr Opin Neurobiol 69: 113–123.

34. Velasco CD, Santarella-Mellwig R, Schorb M, Gao L, Thorn-Seshold O, Llobet A (2023): Microtubule depolymerization contributes to spontaneous neurotransmitter release in vitro. Commun Biol 6: 488.

35. Brodnik ZD, España RA (2015): Dopamine uptake dynamics are preserved under isoflurane anesthesia. Neurosci Lett 606: 129–134.

36. Paxinos G, Watson C (2014): The Rat Brain in Stereotaxic Coordinates, 7th ed. San Diego: Academic Press.

37. Venton BJ, Cao Q (2020): Fundamentals of fast-scan cyclic voltammetry for dopamine detection. Analyst 145: 1158–1168.

38. Venton BJ, Zhang H, Garris PA, Phillips PEM, Sulzer D, Wightman RM (2003): Real-time decoding of dopamine concentration changes in the caudate–putamen during tonic and phasic firing. J Neurochem 87: 1284–1295.

39. Robinson DL, Venton BJ, Heien MLAV, Wightman RM (2003): Detecting Subsecond Dopamine Release with Fast-Scan Cyclic Voltammetry in Vivo. Clin Chem 49: 1763–1773.

40. Bucher ES, Brooks K, Verber MD, Keithley RB, Owesson-White C, Carroll S, et al. (2013): Flexible software platform for fast-scan cyclic voltammetry data acquisition and analysis. Anal Chem 85: 10344–10353.

41. Becker JB, Koob GF (2016): Sex Differences in Animal Models: Focus on Addiction. Pharmacol Rev 68: 242–263.

42. Lynch WJ, Carroll ME (1999): Sex differences in the acquisition of intravenously self-administered cocaine and heroin in rats. Psychopharmacology (Berl) 144: 77–82.

43. Carroll ME, Lynch WJ, Roth ME, Morgan AD, Cosgrove KP (2004): Sex and estrogen influence drug abuse. Trends Pharmacol Sci 25: 273–279.

44. Walker QD, Rooney MB, Wightman RM, Kuhn CM (2000): Dopamine release and uptake are greater in female than male rat striatum as measured by fast cyclic voltammetry. Neuroscience 95: 1061–1070.

45. Calipari ES, Juarez B, Morel C, Walker DM, Cahill ME, Ribeiro E, et al. (2017): Dopaminergic dynamics underlying sex-specific cocaine reward. Nat Commun 8: 13877.

46. Zachry JE, Nolan SO, Brady LJ, Kelly SJ, Siciliano CA, Calipari ES (2021): Sex differences in dopamine release regulation in the striatum. Neuropsychopharmacology 46: 491–499.

47. Becker JB, Chartoff E (2019): Sex differences in neural mechanisms mediating reward and addiction. Neuropsychopharmacology 44: 166–183.

48. Cummings JA, Jagannathan L, Jackson LR, Becker JB (2014): Sex differences in the effects of estradiol in the nucleus accumbens and striatum on the response to cocaine: Neurochemistry and behavior. Drug Alcohol Depend 135: 22–28.

49. Yoest KE, Cummings JA, Becker JB (2014): Estradiol, Dopamine and Motivation. Cent Nerv Syst Agents Med Chem 14: 83–89.

50. Shansky RM (2019): Are hormones a “female problem” for animal research? Science 364: 825–826.

51. Weiss F, Markou A, Lorang MT, Koob GF (1992): Basal extracellular dopamine levels in the nucleus accumbens are decreased during cocaine withdrawal after unlimited-access self-administration. Brain Res 593: 314–318.

52. Schultz W (2016): Dopamine reward prediction-error signalling: a two-component response. Nat Rev Neurosci 17: 183–195.

53. Schultz W, Dayan P, Montague PR (1997): A Neural Substrate of Prediction and Reward. Science 275: 1593–1599.

54. Tsai H-C, Zhang F, Adamantidis A, Stuber GD, Bonci A, de Lecea L, Deisseroth K (2009): Phasic Firing in Dopaminergic Neurons Is Sufficient for Behavioral Conditioning. Science 324: 1080–1084.

55. Grace AA, Bunney BS (1984): The control of firing pattern in nigral dopamine neurons: burst firing. J Neurosci 4: 2877–2890.

56. Olaitan GO, Okojie AK, Paget-Brown R, Dey P, Gatti M, Venton BJ, Lynch WJ (in preparation): Sex-divergent accumbal excitatory-contact remodeling during fentanyl withdrawal. Manuscript in preparation.

57. Li R, Shen Y (2013): An old method facing a new challenge: Re-visiting housekeeping proteins as internal reference control for neuroscience research. Life Sci 92: 747–751.

58. Eaton SL, Roche SL, Llavero Hurtado M, Oldknow KJ, Farquharson C, Gillingwater TH, Wishart TM (2013): Total Protein Analysis as a Reliable Loading Control for Quantitative Fluorescent Western Blotting. PLoS One 8: e72457.

59. Tejeda HA, Bonci A (2019): Dynorphin/kappa-opioid receptor control of dopamine dynamics: Implications for negative affective states and psychiatric disorders. Brain Res 1713: 91–101.

60. Ford CP (2014): The role of D2-autoreceptors in regulating dopamine neuron activity and transmission. Neuroscience 282: 13–22.

