## Supplementary material for "Sex-Dependent Temporal Suppression of Accumbal Dopamine Release During Fentanyl Withdrawal": Main Manuscript

**Supplementary Methods: Subjects**

Rats had ad libitum access to water and food (Teklad LM-485, 7912) and were maintained on a 12-h light/dark cycle (lights on at 7 am). Rats were weighed three times per week, bedding and water bottles were changed twice per week, and health was monitored daily throughout the study. Fentanyl hydrochloride (National Institute on Drug Abuse, Research Triangle Park, NC) was dissolved in sterile saline at 6.25 μg/mL and sterile filtered (0.22 μm; Millipore) before use. Saline controls underwent dopamine assessment and/or tissue collection at time points corresponding to the IntA, EW, and LW conditions, whereas drug-naïve controls underwent these procedures after habituation to the laboratory. Intake analyses include 61 rats (Supplementary Table S5). O457F has 3 recorded IntA days; these were used as recorded in the mixed models, and its 10-day total (mean daily intake × 10) was used only in the Welch test of total infusions. O377M, O387M and O457F contribute intake only (their dopamine values are omitted in the data file and they have no Western data). Missing sessions (35 of 603 rat-days) were filled with the mean of neighboring sessions, or of the rat's other sessions when the first or last sessions were missing; each filled value is listed in the intake data file. Excluding O377M, O387M and O457F did not change the conclusions (Treatment $F_{1,54}=46.93$, $p<0.001$; Sex $F_{1,46}=8.32$, $p=0.006$).

**Supplementary Methods: Western blot procedure**

NAc core tissue was collected 10 min after the end of the voltammetry session, micro-punched on ice, snap-frozen, and stored at −80 °C. Tissue was homogenized in cold RIPA buffer supplemented with protease and phosphatase inhibitor cocktails at the manufacturer-recommended dilution; 20–30 μg protein/lane was resolved on NuPAGE 4–12% gels, transferred to PVDF, and stained with Revert 700 Total Protein Stain (LI-COR). Membranes were incubated overnight at 4 °C with antibodies against DAT (MilliporeSigma MAB369, 1:1000) phospho-DARPP-32 (Thr34) (Cell Signaling Technology 12438, clone D27A4, 1:1000), GABA-A α1R (Synaptic Systems 224-203, 1:1000), GAPDH (Cell Signaling Technology 5174, 1:5000), and β-tubulin (Cell Signaling Technology 2128, 1:1000), together with the other targets reported in the companion study, followed by IRDye 680RD and 800CW secondary antibodies (LI-COR, 1:15,000), and imaged on an Odyssey CLx infrared imaging system (LI-COR). Each band was divided by the total-protein signal of its lane.

Western blot signals were normalized to total lane protein and log10-transformed before analysis because transformation improved residual normality and variance homogeneity for most markers (Supplementary Table S7). Effects of Group (Saline, IntA, EW, LW), Sex, and their interaction were evaluated using two-way ANOVA. Values were expressed as a percentage of the same-sex saline mean (% of control), and the two-way ANOVA was run on log10 of these values; the Group × Sex interaction was re-tested with membrane (a block of lanes from one sex with its own saline lanes) as a fixed factor and as a random intercept (likelihood-ratio test; Supplementary Table S7). Baseline sex differences were tested in saline rats with two-sample t-tests on log10 total-protein-normalized signal. Planned within-sex comparisons used Dunnett-adjusted tests in two families (each fentanyl group versus saline; EW and LW versus IntA), and females and males were compared within each fentanyl group with Holm-adjusted contrasts from the cell-means model (Supplementary Table S9). All tests were two-sided, with significance set at $p < 0.05$. Figures show arithmetic means ± SEM.

**Supplementary Methods: Voltammetry electrodes and recording**

Carbon-fiber microelectrodes were made by aspirating single 7-μm T-650 carbon fibers (Cytec) into glass capillaries (A-M Systems, 615000), pulling them on a vertical puller (PE-22, Narishige), and trimming the exposed fiber to 150–200 μm from the glass seal. Voltammograms were collected with a ChemClamp potentiostat (Dagan) and a UNC breakout box with a 1-MΩ headstage, using an Ag/AgCl wire reference electrode and a 3-kHz low-pass filter, and data were acquired and analyzed with HDCV software (University of North Carolina at Chapel Hill). After each experiment, electrodes were calibrated by flow-injection analysis with dopamine standards in phosphate-buffered saline, and current was converted to concentration assuming a linear response. For the kinetic analysis of the 60-pulse, 60-Hz response, current-versus-time traces were baseline-corrected, smoothed (Savitzky–Golay filter, 5-point window, second-order polynomial), and normalized to peak current. Traces were included if the peak exceeded five times baseline noise, the voltammogram identified dopamine (oxidation peak 0.40–1.00 V), and the signal fell to 50% of peak within 3 s. In vivo, the dopamine oxidation peak appeared at more positive potentials than in vitro (median +0.75 V, interquartile range +0.72 to +0.87 V across included traces) because of Ag/AgCl reference drift and tissue resistance; dopamine was identified by the shape of the background-subtracted voltammogram compared with a dopamine template, and inclusion required an oxidation peak between +0.40 and +1.00 V. The correlation of each trace with the template voltammogram is listed in the kinetics data file; no correlation cutoff was applied. Vmax was fitted to the decay of each calibrated response. Because Km was fixed at 0.2 μM rather than fitted, Vmax and Km are not fully separable, so Vmax is interpreted together with t1/2. These kinetic measures were computed with custom Python code (SciPy) from current–time traces exported from HDCV. Area under the curve was integrated over the actual stimulation values (pulse number, Hz, or μA), and each slope is the least-squares fit of release on those values. Per-rat linear fits had median $R^2$ of 0.92–0.96 in every series; the quadratic term was not significant for the frequency ($p = 0.18$) or current ($p = 0.95$) series, and pulse-number curves bent slightly downward in males ($p = 0.002$) but not in females ($p = 0.23$) (Supplementary Table S6, part B).

**Supplementary Results: Saline versus drug-naïve controls**

First, we compared catheterized saline-administering controls to non-catheterized, drug-naïve rats: the two groups did not form separable clusters in principal-component space across the three stimulation series (Supplementary Figure S1), and linear mixed-effects models run separately in each sex revealed no main or interactive effects of Control Type in any series (Control Type, all $p \geq 0.059$; Control Type × Stimulation Level, all $p \geq 0.27$). With saline-only Controls (n = 5 per sex; drug-naïve rats excluded), female suppression held for frequency AUC (LW vs. Control, $p = 0.0009$), amplitude AUC (Holm $p = 0.007$) and 60-Hz release (Holm $p = 0.010$), and male frequency-AUC suppression held (EW $p = 0.040$; LW $p = 0.047$). Pulse-series contrasts kept their direction and size (female LW −59%; male EW and

LW −46% and −48%) but were not significant after Holm correction, which applied because the Condition × Sex interaction was significant in this subset (e.g., male pulse AUC, EW vs. Control, Holm p = 0.081, uncorrected p = 0.014), as expected with 5 rather than 9 Controls.

**Supplementary Results: Prespecified stimulation ranges and single levels**

Analysis of prespecified pulse intervals further substantiated this loss of release capacity across stimulation demands (Supplementary Table S6). For the low pulse average (6–12 pulses; Figure 2E), release showed significant main effects of Condition and Sex (Condition: $F_{3,54}$=15.73, $p$=1.79×10$^{-7}$, $\eta_p^2$=0.466; Sex: $F_{1,54}$=12.04, $p$=0.0010, $\eta_p^2$=0.182; Condition × Sex: $F_{3,54}$=1.95, $p$=0.1328, $\eta_p^2$=0.098). Females exceeded males under IntA ($p$=0.0028). Within females, release differed between conditions (Control vs. EW: $p$=0.0070; Control vs. LW: $p$=3.25×10$^{-4}$; IntA vs. EW: $p$=0.0038; IntA vs. LW: $p$=2.09×10$^{-4}$; EW vs. LW: $p$=0.0315). Within males, release differed between conditions (Control vs. EW: $p$=0.0299; Control vs. LW: $p$=0.0152; IntA vs. LW: $p$=0.0226), with a trend for IntA vs. EW ($p$=0.0953).

At the intermediate 24-pulse stimulus (Figure 2F), release showed significant main effects of Condition and Sex (Condition: $F_{3,54}$=8.78, $p$=7.73×10$^{-5}$, $\eta_p^2$=0.328; Sex: $F_{1,54}$=14.50, $p$=3.60×10$^{-4}$, $\eta_p^2$=0.212; Condition × Sex: $F_{3,54}$=1.92, $p$=0.1376, $\eta_p^2$=0.096). Females exceeded males under IntA ($p$=0.0018). Within females, release differed between conditions (Control vs. LW: $p$=0.0168; IntA vs. EW: $p$=0.0050; IntA vs. LW: $p$=7.49×10$^{-4}$), with a trend for Control vs. EW ($p$=0.0557). Within males, release differed between conditions (Control vs. EW: $p$=0.0146; Control vs. LW: $p$=0.0071).

For the high pulse average (48–60 pulses; Figure 2G), release showed significant main effects of Condition and Sex (Condition: $F_{3,54}$=12.21, $p$=3.29×10$^{-6}$, $\eta_p^2$=0.404; Sex: $F_{1,54}$=27.24, $p$=2.93×10$^{-6}$, $\eta_p^2$=0.335; Condition × Sex: $F_{3,54}$=2.74, $p$=0.0523, $\eta_p^2$=0.132). Females exceeded males under IntA ($p$=0.0073), early withdrawal ($p$=0.0275), with a trend under Control conditions ($p$=0.0522). Within females, release differed between conditions (Control vs. EW: $p$=0.0315; Control vs. LW: $p$=0.0076; IntA vs. EW: $p$=0.0106; IntA vs. LW: $p$=0.0053). Within males, release differed between conditions (Control vs. EW: $p$=0.0018; Control vs. LW: $p$=0.0060; IntA vs. EW: $p$=0.0450), with a trend for IntA vs. LW ($p$=0.0710).

Evaluation of prespecified frequency bands confirmed that loss of release capacity spanned the frequency spectrum (Figure 3F–H; Supplementary Table S6). For the low frequency average (5–10 Hz; Figure 3F), release showed significant main effects of Condition and Sex (Condition: $F_{3,53}$=6.31, $p$=9.67×10$^{-4}$, $\eta_p^2$=0.263; Sex: $F_{1,53}$=4.35, $p$=0.0418, $\eta_p^2$=0.076; Condition × Sex: $F_{3,53}$=1.01, $p$=0.3969, $\eta_p^2$=0.054). Females exceeded males under Control conditions ($p$=0.0139), early withdrawal ($p$=0.0155). Within females, release differed between conditions (Control vs. EW: $p$=0.0290; Control vs. LW: $p$=7.44×10$^{-5}$; EW vs. LW: $p$=6.47×10$^{-4}$), with a trend for IntA vs. LW ($p$=0.0659). Within males, release differed between conditions (Control vs. LW: $p$=0.0365). For the intermediate 20-Hz stimulus (Figure 3G), release showed significant main effects of Condition and Sex (Condition: $F_{3,53}$=8.92, $p$=6.97×10$^{-5}$, $\eta_p^2$=0.336; Sex: $F_{1,53}$=6.65,

$p$=0.0127, $\eta_p^2$=0.112; Condition × Sex: $F_{3,53}$=1.54, $p$=0.2141, $\eta_p^2$=0.080). Females exceeded males under Control conditions ($p$=0.0416), early withdrawal ($p$=0.0094). Within females, release differed between conditions (Control vs. EW: $p$=0.0090; Control vs. LW: $p$=4.78×10$^{-4}$; EW vs. LW: $p$=0.0365), with a trend for IntA vs. LW ($p$=0.0620). Within males, release differed between conditions (Control vs. EW: $p$=0.0175; Control vs. LW: $p$=0.0459; IntA vs. EW: $p$=0.0498). For the high frequency average (40–60 Hz; Figure 3H), release showed significant main effects of Condition and Sex and a significant Condition × Sex interaction (Condition: $F_{3,53}$=7.72, $p$=2.27×10$^{-4}$, $\eta_p^2$=0.304; Sex: $F_{1,53}$=11.72, $p$=0.0012, $\eta_p^2$=0.181; Condition × Sex: $F_{3,53}$=3.02, $p$=0.0377, $\eta_p^2$=0.146). Female and male values did not differ significantly in any condition (trend under Control conditions, $p_{Holm}$=0.0691, early withdrawal, $p_{Holm}$=0.0691). Within females, release differed between conditions (Control vs. EW: $p_{Holm}$=0.0130; Control vs. LW: $p_{Holm}$=8.64×10$^{-4}$), with a trend for EW vs. LW ($p_{Holm}$=0.0854). Within males, no condition differed significantly (trend for Control vs. EW, $p_{Holm}$=0.0923).

At the maximum burst frequency (60 Hz; Supplementary Figure S2J), release showed significant main effects of Condition and Sex and a significant Condition × Sex interaction (Condition: $F_{3,53}$=6.42, $p$=8.59×10$^{-4}$, $\eta_p^2$=0.267; Sex: $F_{1,53}$=10.49, $p$=0.0021, $\eta_p^2$=0.165; Condition × Sex: $F_{3,53}$=3.43, $p$=0.0235, $\eta_p^2$=0.162). Female and male values did not differ significantly in any condition (trend under Control conditions, $p_{Holm}$=0.0590). Within females, release differed between conditions (Control vs. EW: $p_{Holm}$=0.0086; Control vs. LW: $p_{Holm}$=9.36×10$^{-4}$). Within males, no condition differed significantly.

Analysis of individual current amplitudes detailed this temporal progression (Supplementary Table S6). At the lowest amplitude (100 μA; Figure 4F), release showed significant main effects of Condition and Sex (Condition: $F_{3,53}$=3.38, $p$=0.0248, $\eta_p^2$=0.161; Sex: $F_{1,53}$=5.50, $p$=0.0228, $\eta_p^2$=0.094; Condition × Sex: $F_{3,53}$=0.89, $p$=0.4506, $\eta_p^2$=0.048). Females exceeded males under IntA ($p$=0.0453), early withdrawal ($p$=0.0486). Within females, no condition differed significantly (trend for Control vs. LW, $p$=0.0555). Within males, release differed between conditions (Control vs. EW: $p$=0.0227), with a trend for Control vs. IntA ($p$=0.0894). At the intermediate amplitude (300 μA; Figure 4H), release showed significant main effects of Condition and Sex and a significant Condition × Sex interaction (Condition: $F_{3,54}$=4.53, $p$=0.0066, $\eta_p^2$=0.201; Sex: $F_{1,54}$=14.57, $p$=3.50×10$^{-4}$, $\eta_p^2$=0.213; Condition × Sex: $F_{3,54}$=3.36, $p$=0.0253, $\eta_p^2$=0.157). Female and male values did not differ significantly in any condition (trend under Control conditions, $p_{Holm}$=0.0504, IntA, $p_{Holm}$=0.0770, early withdrawal, $p_{Holm}$=0.0705). Within females, release differed between conditions (Control vs. LW: $p_{Holm}$=0.0256; IntA vs. LW: $p_{Holm}$=0.0296). Within males, no condition differed significantly. At the highest amplitude (500 μA; Figure 4J), release showed significant main effects of Condition and Sex and a significant Condition × Sex interaction (Condition: $F_{3,54}$=9.87, $p$=2.73×10$^{-5}$, $\eta_p^2$=0.354; Sex: $F_{1,54}$=27.71, $p$=2.50×10$^{-6}$, $\eta_p^2$=0.339; Condition × Sex: $F_{3,54}$=4.95, $p$=0.0041, $\eta_p^2$=0.216). Females exceeded males under Control conditions ($p_{Holm}$=0.0156), IntA ($p_{Holm}$=0.0436), early withdrawal ($p_{Holm}$=0.0436). Within females, release differed between conditions (Control vs. LW: $p_{Holm}$=0.0012), with a trend for IntA vs. LW ($p_{Holm}$=0.0500). Within males, release differed between conditions (Control vs. EW: $p_{Holm}$=0.0063).

**Supplementary Results: GABA-A α1R and Western blot sensitivity analyses**

GABA-A α1R showed no significant effect of Group ($F_{3,28} = 2.55$, $p = 0.076$), Sex relative to same-sex saline ($F_{1,28} = 2.45$, $p = 0.129$), or their interaction ($F_{3,28} = 1.71$, $p = 0.187$; Figure 5I).

With membrane as a fixed factor, the GAPDH Group × Sex interaction remained significant ($F_{3,28} = 5.03$, $p = 0.007$), and it was also significant with membrane as a random intercept ($\chi^2_3 = 14.88$, $p = 0.002$); no other marker showed an interaction under either model (all $p \geq 0.06$). With membrane as a fixed factor, the DAT Group effect was significant ($F_{3,26} = 3.47$, $p = 0.030$). Log10 transformation improved residual normality for GAPDH, β-tubulin, phospho-DARPP-32 and GABA-A α1R but not for DAT (Shapiro–Wilk $p = 0.23$ raw and 0.049 log10; Supplementary Table S7). Rank-based (aligned rank transform) ANOVAs gave the same conclusions as the primary analyses for all five proteins, whereas removing the most extreme animal from each Group × Sex cell left the GAPDH interaction ($p = 0.004$) and the β-tubulin sex difference ($p < 0.001$) intact, made the DAT Group effect significant ($p = 0.020$), and produced Group × Sex interactions for phospho-DARPP-32 ($p = 0.024$) and GABA-A α1R ($p < 0.001$; Supplementary Table S7). Excluding the two lanes with the lowest total-protein signal (O391M and O472M; TPN 4.81 and 2.55), the GAPDH Group × Sex interaction remained significant ($F_{3,30} = 3.24$, $p = 0.036$; full data $p = 0.014$).

### Supplementary Figure S1

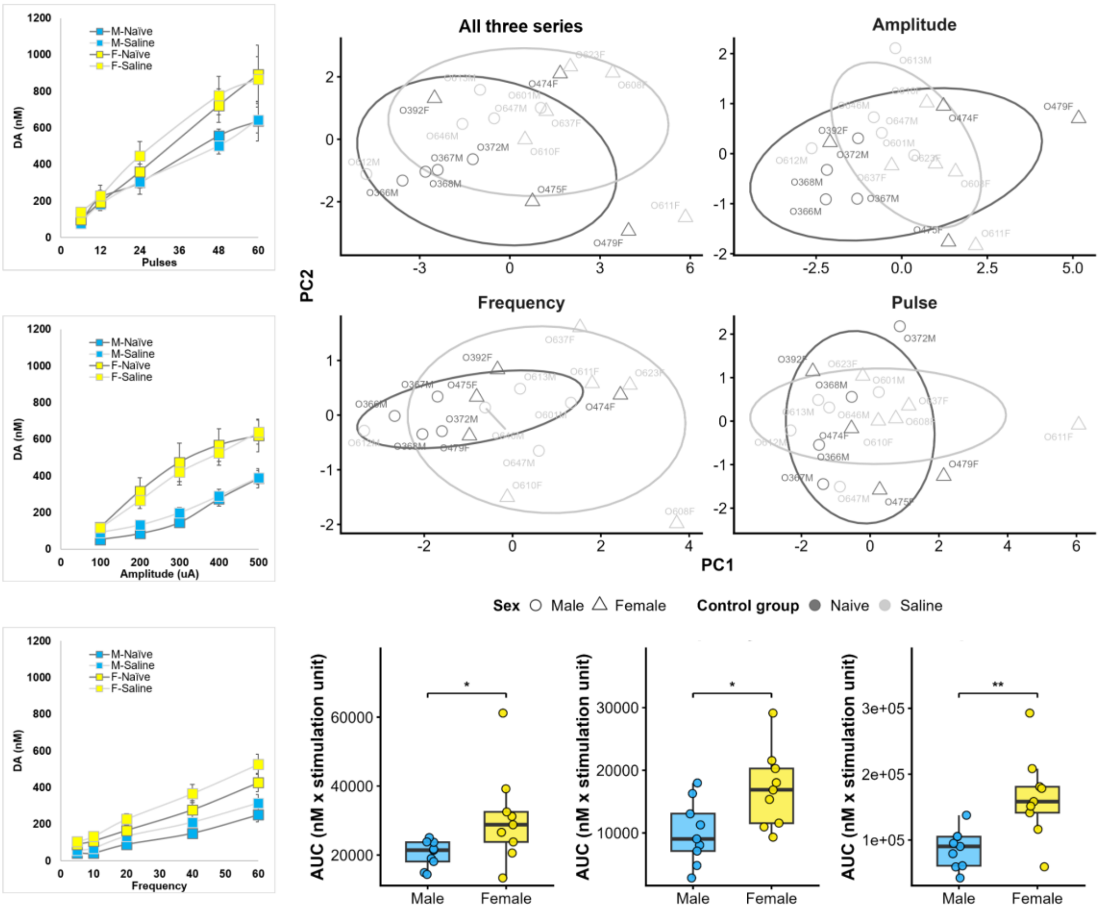


Supplementary Figure S1. Drug-naive and saline Controls do not differ on the voltammetric endpoints. The two control sources do not form separable clusters in principal-component space, and linear mixed-effects models run separately in each sex (Control Type × Stimulation Level, random intercept for animal) showed no effect of Control Type (all p≥0.059) or Control Type × Stimulation Level interaction (all p≥0.27) in any series, justifying consolidation of drug-naive and saline animals into a single Control group for the FSCV analyses. Full statistics are listed in Supplementary Table S1. Left column, pulse, current, and frequency series for naive and saline controls of each sex; centre, principal-component scores for all three series combined and for each series; bottom row, pooled Control AUC for the three series by sex (*p<0.05, **p<0.01, females vs. males). Series data are mean ± SEM; box plots show median and interquartile range; points are individual animals.

### Supplementary Figure S2

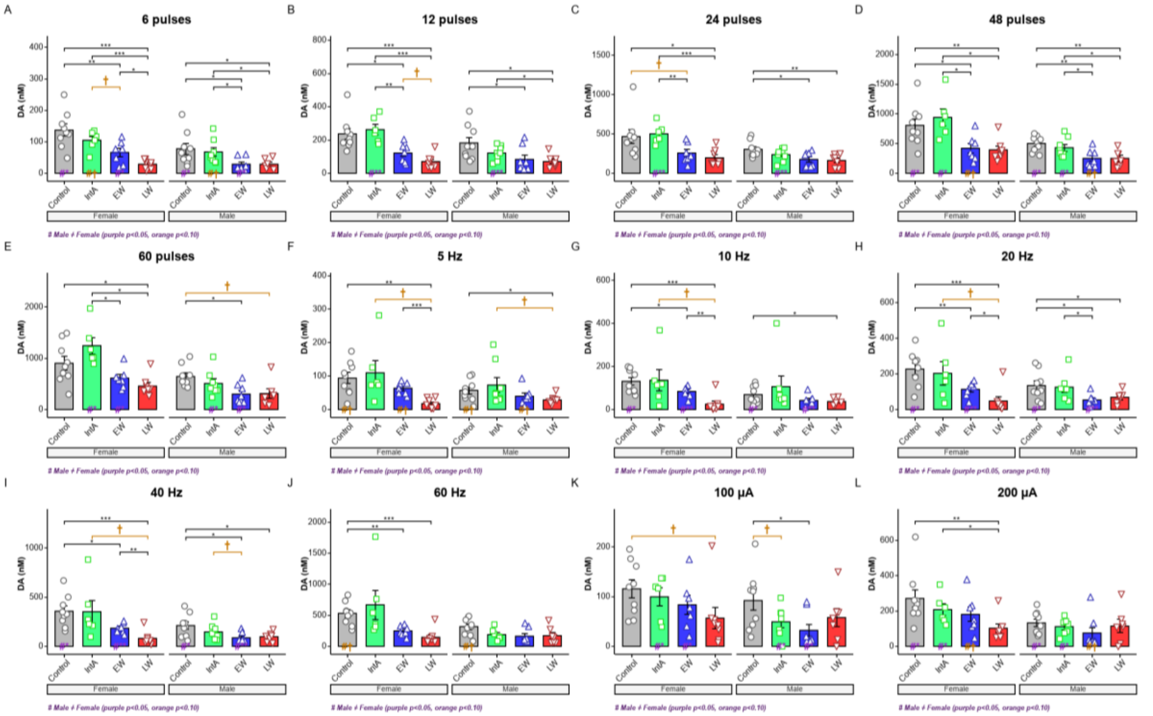


Supplementary Figure S2. Full FSCV atlas. Sex × Condition means ± SEM on the nM scale for every single stimulation level and prespecified range average: (A–E) 6, 12, 24, 48, and 60 pulses; (F–J) 5, 10, 20, 40, and 60 Hz; (K–O) 100, 200, 300, 400, and 500 µA; (P–R) 6–12, 24, and 48–60 pulses; (S–U) 5–10, 20, and 40–60 Hz. Brackets mark within-sex differences between conditions (*$p<0.05$, **$p<0.01$, ***$p<0.001$, ****$p<0.0001$; † $p<0.10$); # marks a female–male difference within a condition (purple $p<0.05$, orange $p<0.10$). Brackets and # marks are Welch tests, Holm-adjusted when the Condition × Sex interaction was significant ($p<0.05$); brackets include comparisons with Control. Points are individual animals.

### Supplementary Figure S2 (continued)

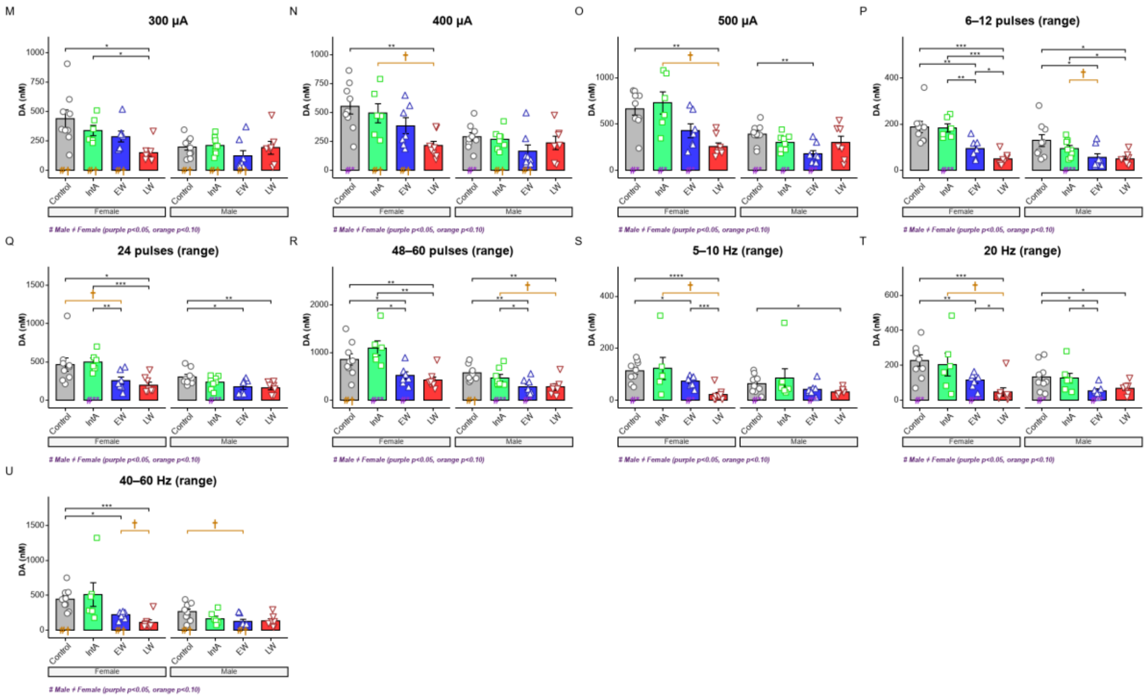


Supplementary Figure S2 (continued). Panels M–U; see the legend above.

### Supplementary Figure S3

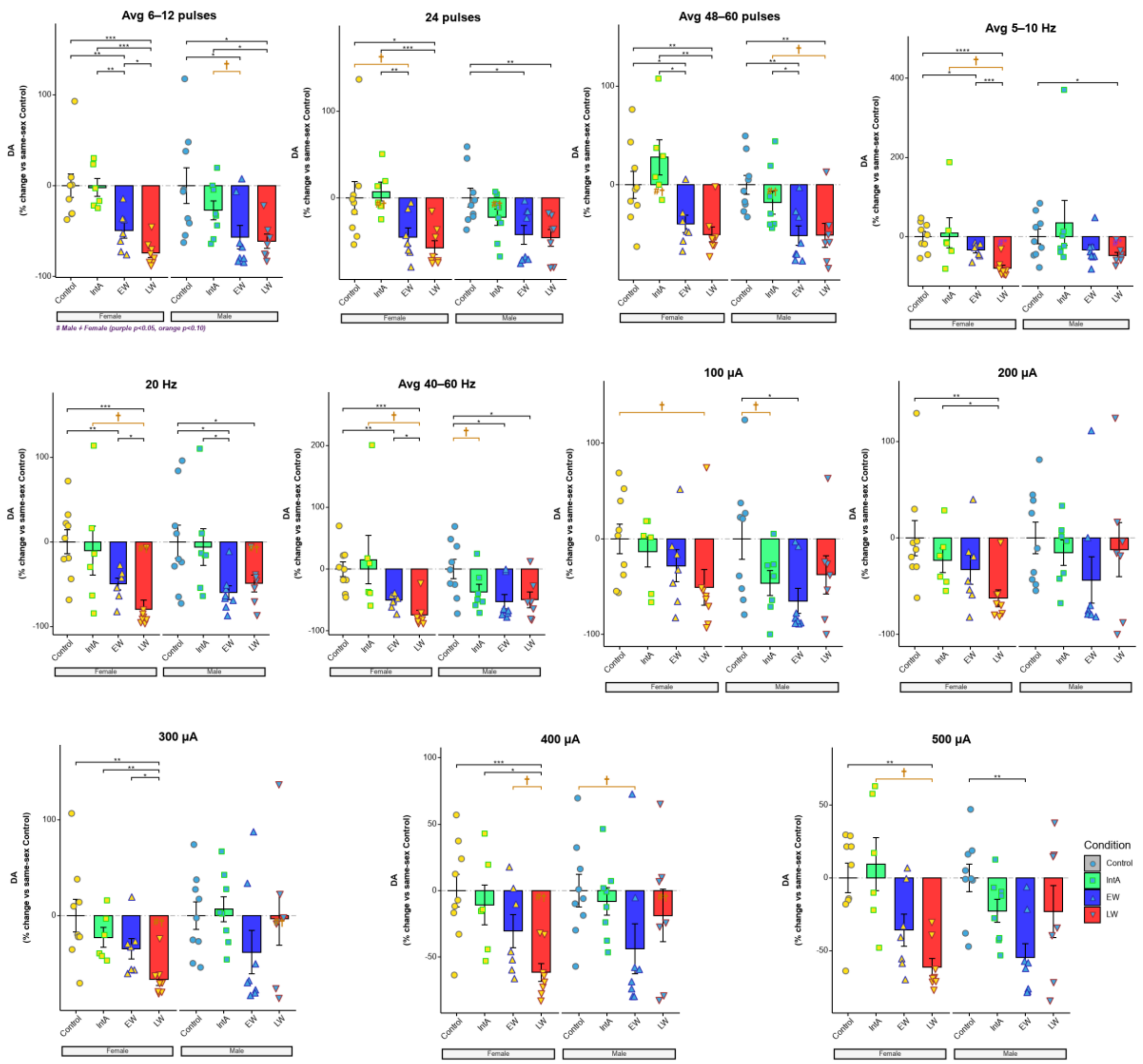


Supplementary Figure S3. Per-animal percentage change from the same-sex pooled Control mean across FSCV dopamine measurements. Percentage change was calculated as 100 x [(individual value/same-sex Control mean) - 1]. Control observations retain individual variability around a mean of 0%. Brackets mark Welch tests between conditions within each sex, including comparisons with Control (IntA, EW and LW versus Control and versus each other), Holm-adjusted when the Condition × Sex interaction was significant (p<0.05); † p<0.10; # marks a female–male difference (purple p<0.05, orange p<0.10).

### Supplementary Figure S4

Rat ID colour key: **Control/Saline** **IntA** **EW** **LW**

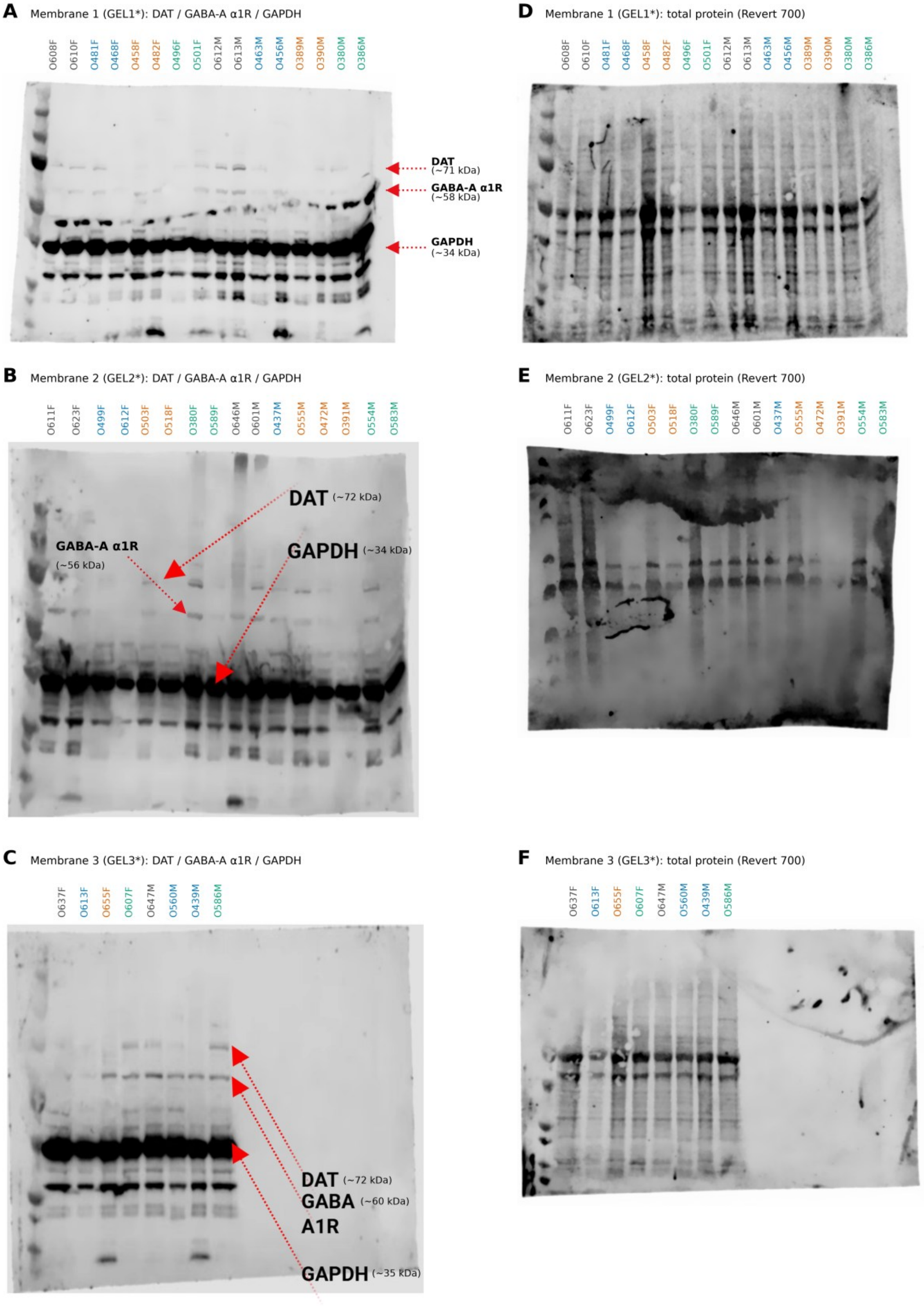

Supplementary Figure S4. Uncropped Western blot membranes for DAT, GABA-A α1R and GAPDH, with the total-protein image of each membrane. (A–C) The three membranes (GEL1, 16 lanes; GEL2, 16 lanes; GEL3, 8 lanes), each shown once, with rat IDs above the lanes (Control/Saline, grey; IntA, blue; EW, orange; LW, green). Arrows mark DAT (~72 kDa), GABA-A α1R (~56–60 kDa) and GAPDH (~34–35 kDa). (D–F) Revert 700 total-protein images of the same membranes, in the same order. These membranes supply the GAPDH, DAT and GABA-A α1R values in Figure 5E, 5G and 5I.

### Supplementary Figure S5

Rat ID colour key: **Control/Saline** **IntA** **EW** **LW**

**A** p-DARPP-32 (Thr34) membrane (GEL 1B)

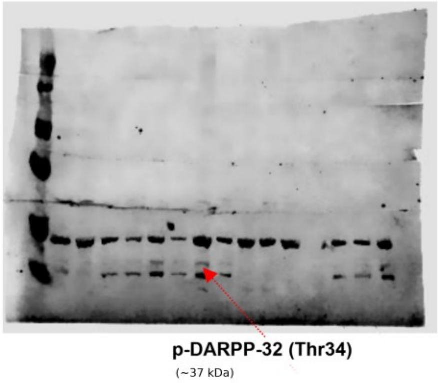


**B** GAD1 membrane

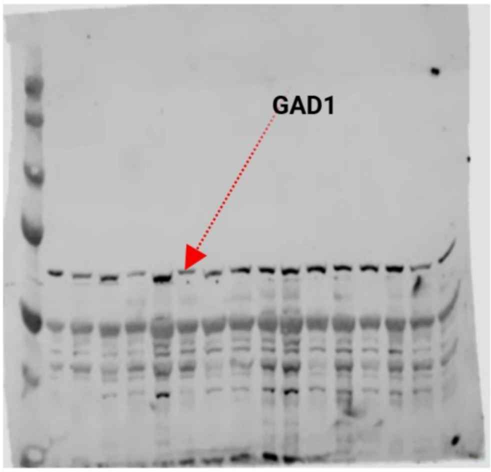


**C** β-tubulin / adenosine A1R membrane (Gel 1B)

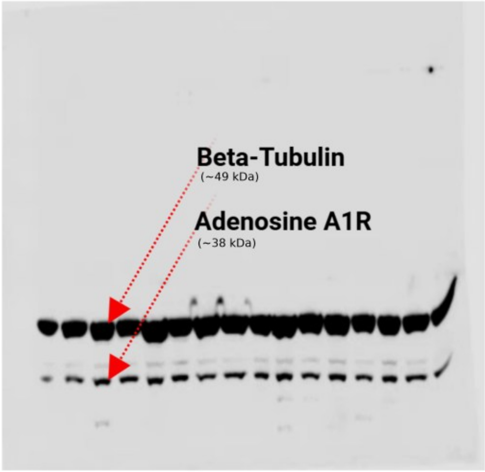


**D** β-tubulin / adenosine A1R membrane (Gel 2)

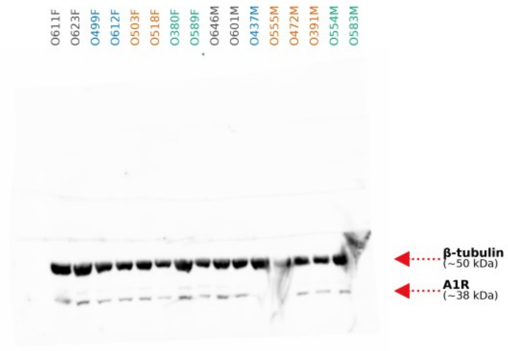


**E** β-tubulin / adenosine A1R membrane (Gel 3)

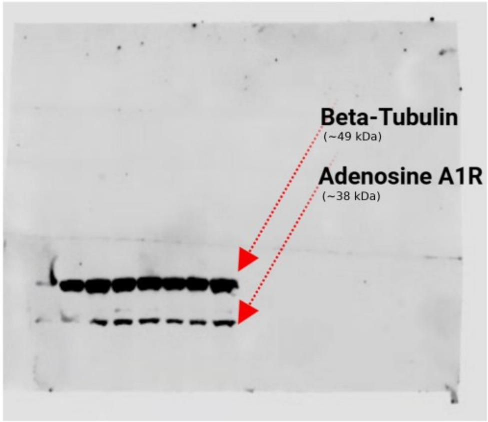

Supplementary Figure S5. Additional uncropped membranes. (A) p-DARPP-32 (Thr34) (~37 kDa). (B) GAD1. (C–E) β-tubulin (~50 kDa) and adenosine A1R (~38 kDa) on three membranes. Rat IDs are shown above the lanes of panel D; colours as in Supplementary Figure S4. Each membrane is shown once. These membranes supply the β-tubulin and p-DARPP-32 values in Figure 5F and 5H; GAD1 and adenosine A1R are reported in the companion study.

### Supplementary Figure S6

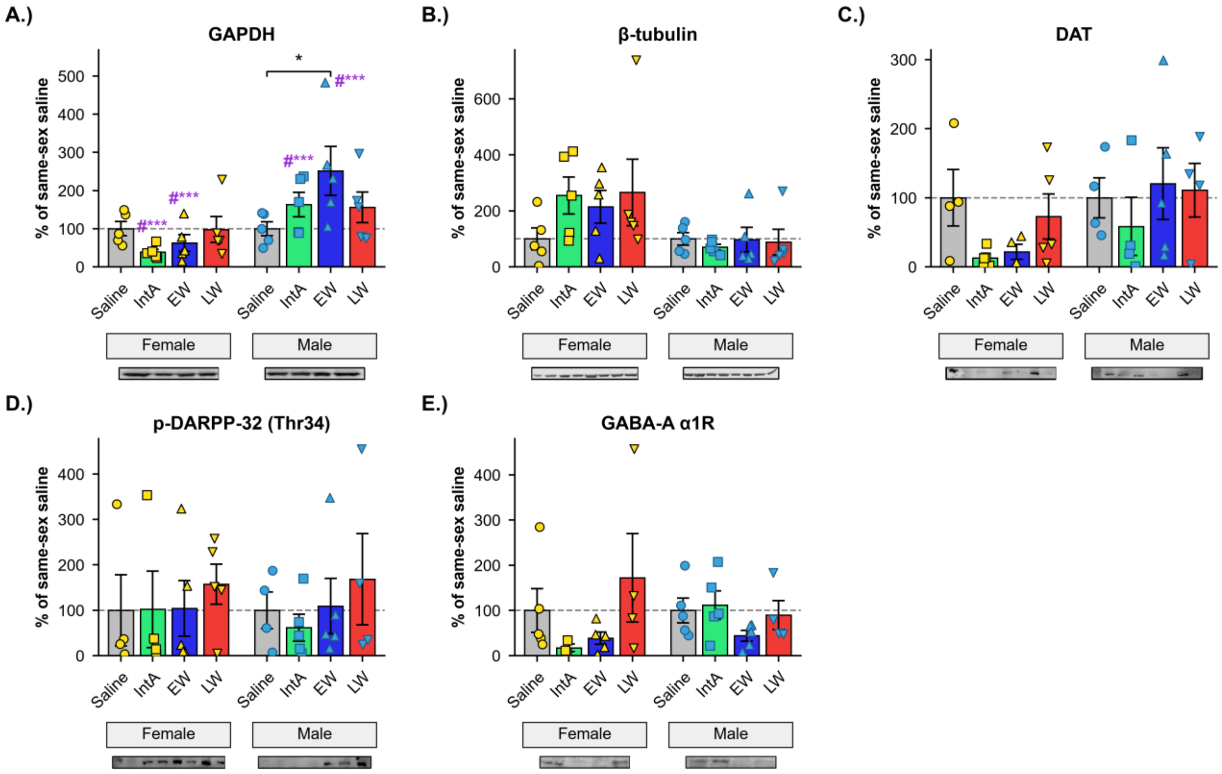


Supplementary Figure S6. Western blot markers expressed as percent of the same-sex saline mean. (A) GAPDH, (B) β-tubulin, (C) total DAT, (D) phospho-DARPP-32 (Thr34) and (E) GABA-A α1R in NAc core tissue, with representative blots below each panel. Values were log10-transformed for analysis and are plotted untransformed. *p<.05 versus same-sex saline (Dunnett; Supplementary Table S9). # marks a female–male difference within a fentanyl group on percent of same-sex saline (Holm-adjusted; #*** p<.001). Saline is 100% in both sexes by construction, so baseline sex differences are shown in Figure 5. Bars are mean ± SEM; points are individual animals.

### Supplementary Tables

**Supplementary Table S1.** Drug-naïve versus saline Controls, analysed separately in each sex. Part A: for each stimulation series, a mixed model of Control Type × Stimulation Level with a random intercept for animal (Wald $\chi^2$, df, p). Part B: for each endpoint, Welch t-test (mean ± SEM, n, t, df, p), Hedges g with 95% CI, and a TOST equivalence test with bounds of ±1 SD (naïve and saline SD pooled within sex).

**Supplementary Table S2.** Per-cell sample sizes (Sex × Condition × assay). Number of animals contributing to each FSCV endpoint and each Western blot endpoint (total DAT, phospho-DARPP-32, GABA-A α1R, GAPDH, and β-tubulin).

Supplementary Table S3. Per-rat clearance half-time, release per pulse, and $V_{max}$ (printed below).

**Supplementary Table S4.** Bayes factors for evoked dopamine release (nM scale). BayesFactor::anovaBF (y ~ Condition × Sex, whichModels = 'withmain') with default JZS priors (rscaleFixed = 'medium', r = 0.5), 100,000 iterations and set.seed(20261027). $BF_{10}$ is given for Condition, Sex and Condition × Sex by model comparison, with matched-model inclusion Bayes factors and Lee and Wagenmakers evidence categories.

**Supplementary Table S5.** Baseline comparisons. Part A: intake equivalence across withdrawal groups. Sex × withdrawal-group and withdrawal-group × day mixed-effects results; overall intake did not differ by sex across the assigned withdrawal groups, though daily intake patterns diverged significantly among the three groups across the 10-day access period prior to voltammetry or protein analyses. Part B: baseline sex differences in Control rats (two-sample t-tests and Hedges g) for every voltammetric endpoint.

**Supplementary Table S6.** Within-sex Condition ANOVAs for every FSCV endpoint (nM scale). One-way ANOVA of Condition (Control, IntA, EW, LW), run separately in females and males, with F, df, p and $\eta^2$ (part A). Part B: linearity of each stimulation series, from per-rat fits on the actual stimulation values (pulse number, Hz, μA): linear $R^2$, added $R^2$ from a quadratic term, and a one-sample test of the standardized quadratic term (t test and Wilcoxon).

Supplementary Table S7. Western blot sensitivity analyses and residual diagnostics (printed below).

**Supplementary Table S8. Two-way ANOVA (Group × Sex; Saline, IntA, EW, LW; Type II) on log10-transformed, total-protein-normalized signal for GAPDH, β-tubulin, total DAT, phospho-DARPP-32 (Thr34), and GABA-A α1R. All effects are tested on log10 of values expressed as percent of the same-sex saline mean.**

**Supplementary Table S9. Part A: mean percent of same-sex saline and mean relative expression (± SEM) for the five Western markers. Part B: for the Western markers, within-sex Dunnett comparisons against saline and against IntA and female–male Holm-adjusted contrasts within each fentanyl group (log10 percent of same-sex**

**saline), and baseline saline t-tests (log10 relative expression); for clearance half-time, [DA]p and Vmax (Figure 5B–D), planned Welch t-tests, uncorrected because no Condition × Sex interaction was significant. The Mark column gives the symbol shown in Figure 5 or Supplementary Figure S6.**

Supplementary Tables S1–S9 are provided as an accompanying Excel workbook, one sheet per table; Supplementary Tables S3 and S7 are also printed below.

### Supplementary Table S3

Supplementary Table S3. Clearance half-time ($t_{1/2}$), release per pulse ($[DA]_p$), and maximal uptake rate ($V_{max}$) for each rat, from the 60-pulse, 60-Hz response (Figure 5). All listed rats met the trace inclusion criteria in the Supplementary Methods (peak above five times baseline noise, dopamine voltammogram with an oxidation peak at 0.40–1.00 V, and decay to 50% of peak within 3 s). Dashes mark rats for which $[DA]_p$ and $V_{max}$ were not estimated because the Michaelis–Menten fit to the decay was poor. Kinetic inclusion was judged on each rat's 60-pulse trace, so kinetic n differ from those in Figures 2–4 (Supplementary Table S2). Control, pooled saline and drug-naïve controls; IntA, immediately after the final intermittent-access session; EW, early withdrawal (days 1–3); LW, late withdrawal (days 13–16).

| Rat | Sex | Condition | $t_{1/2}$ (s) | $[DA]_p$ (nM) | $V_{max}$ (µM/s) |
|---|---|---|---|---|---|
| 0475F | Female | Control | 0.741 | 24.0 | 0.42 |
| 0479F | Female | Control | 0.456 | 47.7 | 1.66 |
| 0610F | Female | Control | 0.516 | 19.8 | 0.86 |
| 0611F | Female | Control | 0.339 | 21.5 | 1.67 |
| 0623F | Female | Control | 0.636 | 18.6 | 0.61 |
| 0367M | Male | Control | 0.673 | 26.0 | 0.82 |
| 0368M | Male | Control | 0.447 | 12.2 | 0.45 |
| 0458M | Male | Control | 0.964 | 22.2 | 0.45 |
| 0601M | Male | Control | 2.083 | 35.5 | 0.40 |
| 0612M | Male | Control | 0.466 | — | — |
| 0613M | Male | Control | 1.148 | 8.2 | 0.27 |
| 0646M | Male | Control | 0.516 | — | — |
| 0647M | Male | Control | 0.441 | — | — |
| 0468F | Female | IntA | 0.770 | 51.2 | 1.21 |
| 0481F | Female | IntA | 0.474 | 35.9 | 1.10 |
| 0499F | Female | IntA | 0.413 | — | — |
| 0612F | Female | IntA | 0.416 | 30.9 | 1.34 |
| 0613F | Female | IntA | 0.149 | 41.9 | 3.87 |
| 0388M | Male | IntA | 0.483 | — | — |
| 0439M | Male | IntA | 0.545 | 13.4 | 0.50 |
| 0453M | Male | IntA | 0.293 | 33.2 | 1.54 |
| 0456M | Male | IntA | 0.979 | — | — |
| 0463M | Male | IntA | 0.525 | 9.0 | 0.39 |
| 0560M | Male | IntA | 0.568 | — | — |
| 0458F | Female | EW | 0.487 | 16.6 | 0.73 |
| 0467F | Female | EW | 0.533 | 26.0 | 0.63 |
| 0482F | Female | EW | 0.655 | — | — |
| 0490F | Female | EW | 1.376 | 7.1 | 0.15 |
| 0503F | Female | EW | 1.540 | 13.3 | 0.21 |
| 0655F | Female | EW | 0.517 | — | — |
| 0395M | Male | EW | 1.017 | 3.7 | 0.13 |
| 0414M | Male | EW | 0.393 | 8.0 | 0.56 |
| 0555M | Male | EW | 0.812 | 13.2 | 0.44 |
| 0459F | Female | LW | 1.776 | 9.3 | 0.15 |
| 0496F | Female | LW | 0.439 | 8.4 | 0.57 |
| 0501F | Female | LW | 0.508 | 10.3 | 0.54 |
| 0502F | Female | LW | 0.472 | 10.8 | 0.71 |
| 0589F | Female | LW | 0.607 | 6.5 | 0.31 |
| 0607F | Female | LW | 0.794 | 7.3 | 0.24 |
| 0375M | Male | LW | 0.617 | 5.2 | 0.28 |
| 0376M | Male | LW | 0.783 | 2.4 | 0.25 |
| 0380M | Male | LW | 0.531 | 2.1 | 0.14 |

| | | | | | |
|---|---|---|---|---|---|
| O381M | Male | LW | 0.555 | 1.2 | 0.33 |
| O386M | Male | LW | 2.197 | 9.9 | 0.18 |
| O438M | Male | LW | 0.450 | 5.7 | 0.31 |
| O554M | Male | LW | 0.552 | 6.3 | 0.26 |
| O583M | Male | LW | 0.737 | 38.2 | 0.77 |

### Supplementary Table S7

Supplementary Table S7. Western blot sensitivity analyses. Group × Sex interaction on log10 signal without and with membrane as a fixed factor or a random intercept (likelihood-ratio test), the Group effect with membrane as a fixed factor, and Shapiro–Wilk (residual normality) and Brown–Forsythe (variance homogeneity) p values for raw versus log10-transformed signal. The last two columns give the Group × Sex interaction from a rank-based (aligned rank transform) ANOVA and from the log10 ANOVA after removing the most extreme animal (largest absolute deviation from the cell median) from each Group × Sex cell.

| **Marker** | **N (membranes)** | **Group × Sex, log10** | **+ membrane (fixed)** | **+ membrane (random, LRT)** | ***Group, + membrane (fixed)*** | ***Shapiro–Wilk p, raw / log10*** | ***Brown–Forsythe p, raw / log10*** | ***Membrane variance share*** | ***Group × Sex, rank-based (ART)*** |
|---|---|---|---|---|---|---|---|---|---|
| GAPDH | 40 (6) | $F_{3,32}=4.15$, $p=0.014$ | $F_{3,28}=5.03$, $p=0.007$ | $\chi^2_3=14.88$, $p=0.002$ | $F_{3,31}=0.55$, $p=0.654$ | 0.009 / 0.937 | 0.767 / 0.914 | 0.16 | $F_{3,32}=4.32$, $p=0.011$ |
| β-tubulin | 40 (6) | $F_{3,32}=1.93$, $p=0.144$ | $F_{3,28}=2.26$, $p=0.104$ | $\chi^2_3=7.25$, $p=0.064$ | $F_{3,31}=0.67$, $p=0.579$ | 0.001 / 0.212 | 0.924 / 0.845 | 0.07 | $F_{3,32}=1.68$, $p=0.190$ |
| DAT | 35 (6) | $F_{3,27}=0.71$, $p=0.553$ | $F_{3,23}=0.47$, $p=0.705$ | $\chi^2_3=2.49$, $p=0.477$ | $F_{3,26}=3.47$, $p=0.030$ | 0.231 / 0.049 | 0.555 / 0.879 | 0.12 | $F_{3,27}=0.54$, $p=0.656$ |
| p-DARPP-32 (Thr34) | 36 (6) | $F_{3,28}=0.07$, $p=0.974$ | $F_{3,24}=0.01$, $p=0.999$ | $\chi^2_3=0.28$, $p=0.963$ | $F_{3,24}=0.79$, $p=0.511$ | 0.001 / 0.748 | 0.957 / 0.987 | 0.00 | $F_{3,28}=0.10$[illegible] $p=0.962$ |
| GABA-A α1 | 36 (6) | $F_{3,28}=1.71$, $p=0.187$ | $F_{3,24}=1.37$, $p=0.275$ | $\chi^2_3=6.01$, $p=0.111$ | $F_{3,27}=2.30$, $p=0.100$ | 0.000 / 0.576 | 0.367 / 0.949 | 0.04 | $F_{3,28}=1.74$[illegible] $p=0.182$ |
